# ZccE is a Novel P-type ATPase That Protects *Streptococcus mutans* Against Zinc Intoxication

**DOI:** 10.1101/2022.03.29.486188

**Authors:** Tridib Ganguly, Alexandra Peterson, Marissa Burkholder, Jessica K. Kajfasz, Jacqueline Abranches, José A. Lemos

## Abstract

Zinc is a trace metal that is essential to all forms of life, but that becomes toxic at high concentrations. Because it has both antimicrobial and anti-inflammatory properties and low toxicity to mammalian cells, zinc has been used as a therapeutic agent for centuries to treat a variety of infectious and non-infectious conditions. While the usefulness of zinc-based therapies in caries prevention is controversial, zinc is incorporated into toothpaste and mouthwash formulations to prevent gingivitis and halitosis. Despite this widespread use of zinc in oral healthcare, the mechanisms that allow *Streptococcus mutans*, a keystone pathogen in dental caries and prevalent etiological agent of infective endocarditis, to overcome zinc toxicity are largely unknown. Here, we discovered that *S. mutans* is inherently more tolerant to high zinc stress than all other species of streptococci tested, including commensal streptococci associated with oral health. Using a transcriptome approach, we uncovered several potential strategies utilized by *S. mutans* to overcome zinc toxicity. Among them, we identified a previously uncharacterized P-type ATPase transporter and cognate transcriptional regulator, which we named ZccE and ZccR respectively, as responsible for the remarkable high zinc tolerance of *S. mutans*. In addition to zinc, we found that ZccE, which was found to be unique to *S. mutans* strains, mediates tolerance to at least three additional metal ions, namely cadmium, cobalt, and copper. Loss of the ability to maintain zinc homeostasis when exposed to high zinc stress severely disturbed Zinc:Manganese ratios, leading to heightened peroxide sensitivity that was alleviated by manganese supplementation. Finally, we showed that the ability of the Δ*zccE* strain to stably colonize the rat tooth surface after topical zinc treatment was significantly impaired, providing proof of concept that ZccE and ZccR are suitable targets for the development of antimicrobial therapies specifically tailored to kill *S. mutans*.

**Author summary:** Dental caries is an overlooked infectious disease affecting more than 50% of the adult population. While several bacteria that reside in dental plaque have been associated with caries development and progression, *Streptococcus mutans* is deemed a keystone caries pathogen due to its capacity to modify the dental plaque environment in a way that is conducive with disease development. Zinc is an essential trace metal to life but very toxic when encountered at high concentrations, to the point that it has been used as an antimicrobial for centuries. Despite the widespread use of zinc in oral healthcare products, little is known about the mechanisms utilized by oral bacteria to overcome its toxic effects. In this study, we discovered that *S. mutans* can tolerate exposure to much higher levels of zinc than closely related streptococcal species, including species that antagonize *S. mutans* and are associated with oral health. In this study, we identified a new metal transporter, named ZccE, as directly responsible for the inherently high zinc tolerance of *S. mutans*. Because ZccE is not present in other bacteria, our findings provide a new target for the development of a zinc-based therapy specifically tailored to kill *S. mutans*.

## Introduction

The first row *d*-block metal ions cobalt (Co), copper (Cu), iron (Fe), manganese (Mn), nickel (Ni) and zinc (Zn) are essential trace metals due to their roles in intracellular signaling and incorporation to structural and catalytic domains of proteins that perform the biological processes required for life (1, 2). On the flip side, these metals are toxic when present in excess, such that the ability to maintain metal homeostasis is absolutely critical for the survival of all forms of life (1–3). To prevent or fight infections, the host activates a number of metal ion withholding strategies, collectively termed nutritional immunity, that ultimately starve pathogens of essential biometals such as Fe, Mn, and Zn (4). For example, Fe in host fluids and tissues is sequestered by transferrin and lactoferrin, or stored within hepatocytes (reviewed in (5)). Both Mn and Zn are primarily sequestered by calprotectin, a heterodimer of the S100A8 and S100A9 proteins produced and secreted in large quantities by neutrophils (reviewed in (6)). At the same time, the host is capable of harnessing metal ions to intoxicate invading pathogens. For example, Cu and Zn are released in large quantities within phagocytic cells to kill invading pathogens (3, 7).

Among the most biologically relevant trace metals, Zn is unique because it does not undergo redox-cycling and, therefore, should not be classified as a transition metal as is often the case. Due to its top position in the Irving-Williams series, Zn can bind more avidly and form more stable interactions with metal ligands when compared to most other metal ions. As a result, Zn is poisonous when in excess due to adventitious binding to non-cognate metalloproteins or even to residues that should not be occupied by metals, a process known as protein mismetallation, (7, 8). Moreover, Zn is critical for proper functioning of the immune system as it is required for activation of pro-inflammatory signaling pathways, neutrophil extracellular trap (NET) formation and, as indicated above, it can be mobilized into phagosomes to kill invading pathogens (3, 4, 9). Because it has antimicrobial properties, low toxicity to mammalian cells, and stimulates immune cell function, Zn has long been used as a therapeutic agent to treat or prevent a variety of medical conditions, from wound healing to gingivitis, or as an adjuvant in the treatment of common colds and other viral infections.

While the significance of Zn deprivation in host-pathogen interactions and the mechanisms utilized by bacteria to overcome Zn deprivation have been studied to some depth (reviewed in (8, 10), the physiological consequences of Zn poisoning and the mechanisms by which pathogens overcome Zn intoxication are less well understood. In the initial response to toxic levels of Zn, bacteria have been shown to induce the expression and activity of membrane-associated Zn efflux systems to remove Zn from their cytosol. To date, three types of Zn efflux systems have been described in bacteria, namely P-type ATPases, resistance <u>n</u>odulation <u>d</u>ivision (RND) pumps, and <u>c</u>ation <u>d</u>iffusion facilitator (CDF) metal/H^+^ antiporters (11). In *Escherichia coli* (the Gram-negative paradigm), the P-type ATPase ZntA is the major Zn efflux pump, whereas *Bacillus subtilis* (the Gram-positive paradigm) has been shown to utilize both a P-type ATPase (known as CadA) and CDF antiporter (known as CzcD) to pump Zn out of the cell (12–15). In all streptococci studied to date, a CDF-type antiporter, homologous to the *B. subtilis* CzcD, has been shown to serve as the major Zn-resistance determinant, with several reports showing that inactivation of *czcD* significantly diminishes bacterial virulence (9, 16–22).

A resident of dental biofilms, *Streptococcus mutans* is a keystone pathogen of dental caries, one of the most prevalent and overlooked infectious diseases in the world (23, 24). In addition to its well-recognized role as a cariogenic pathogen, *S. mutans* is also implicated in infective endocarditis (25), with a recent retrospective study associating transient *S. mutans* bloodstream infections (BSI) with the highest risk of developing into infective endocarditis when compared to other streptococcal BSI (26). In oral health, Zn has been incorporated into mouthwash and toothpaste formulations to prevent and treat halitosis and gingivitis (27, 28). In dental caries, the usefulness of Zn as a therapeutic agent is controversial, with a number of studies concluding that Zn is not effective contrasting with studies lauding Zn as a highly effective anti-plaque or anti-caries agent (29–33). Controversy aside, mechanistic studies have shown that high concentrations of Zn inhibits the activity of glycolytic enzymes of oral streptococci *in vitro* and acid production by dental plaque *in vivo* (33–36). Moreover, Zn and fluoride were shown to act synergistically, impairing *S. mutans’* acid tolerance as well as its ability to synthesize extracellular polymers of glucan from dietary sucrose (37).

Despite the widespread incorporation of Zn to oral healthcare products, very little is known about the mechanisms utilized by oral bacteria, *S. mutans* included, to maintain Zn homeostasis. Recently, we showed that inactivation of *adcABC*, which codes for an ABC-type Zn-specific transporter, drastically reduced the ability of *S. mutans* to colonize the dental biofilm in a rat model, indicating that host-mediated mechanisms for Zn sequestration, competition with microbial commensals, or both, restrict Zn availability to *S. mutans* in dental plaque (38). In this report, we sought to uncover the mechanisms utilized by *S. mutans* to cope with toxic levels of Zn. We discovered that *S. mutans* exhibits much higher tolerance to Zn than all other streptococci tested, including several commensal oral streptococci associated with oral health. Through transcriptome and bioinformatic analyses, we identified a previously uncharacterized P-type ATPase transporter and cognate MerR-type transcriptional regulator as directly responsible for the remarkable tolerance of *S. mutans* against Zn intoxication. *In vitro* and *in vivo* characterizations of a strain lacking this novel Zn efflux system and the fact that it is unique to *S. mutans* indicates that it is a potential target for the development of Zn-based antimicrobial therapies specifically tailored to kill *S. mutans*.

## Results

### *S. mutans* is inherently more tolerant to Zn intoxication when compared to other streptococci

To uncover the mechanisms utilized by *S. mutans* to cope with toxic levels of Zn, we first determined the minimum inhibitory concentration (MIC) of ZnSO_4_ in a panel of 17 streptococcal strains from 9 different species that were in our lab collection. The panel included *S. mutans* and *S. sobrinus* (caries-associated streptococci), oral commensals (e.g., *S. gordonii, S. sanguinis*), and pyogenic streptococci *S. agalactiae* and *S. pyogenes*. Remarkably, all *S. mutans* strains displayed much higher Zn tolerance (MIC = 4 mM) when compared to the other species tested (MIC ranging from 0.5 to 1 mM) (**Table 1**). The intrinsically high Zn tolerance of the *S. mutans* strains when compared to other streptococci was further validated in disc diffusion assays (**Fig 1**).

**Fig 1.**
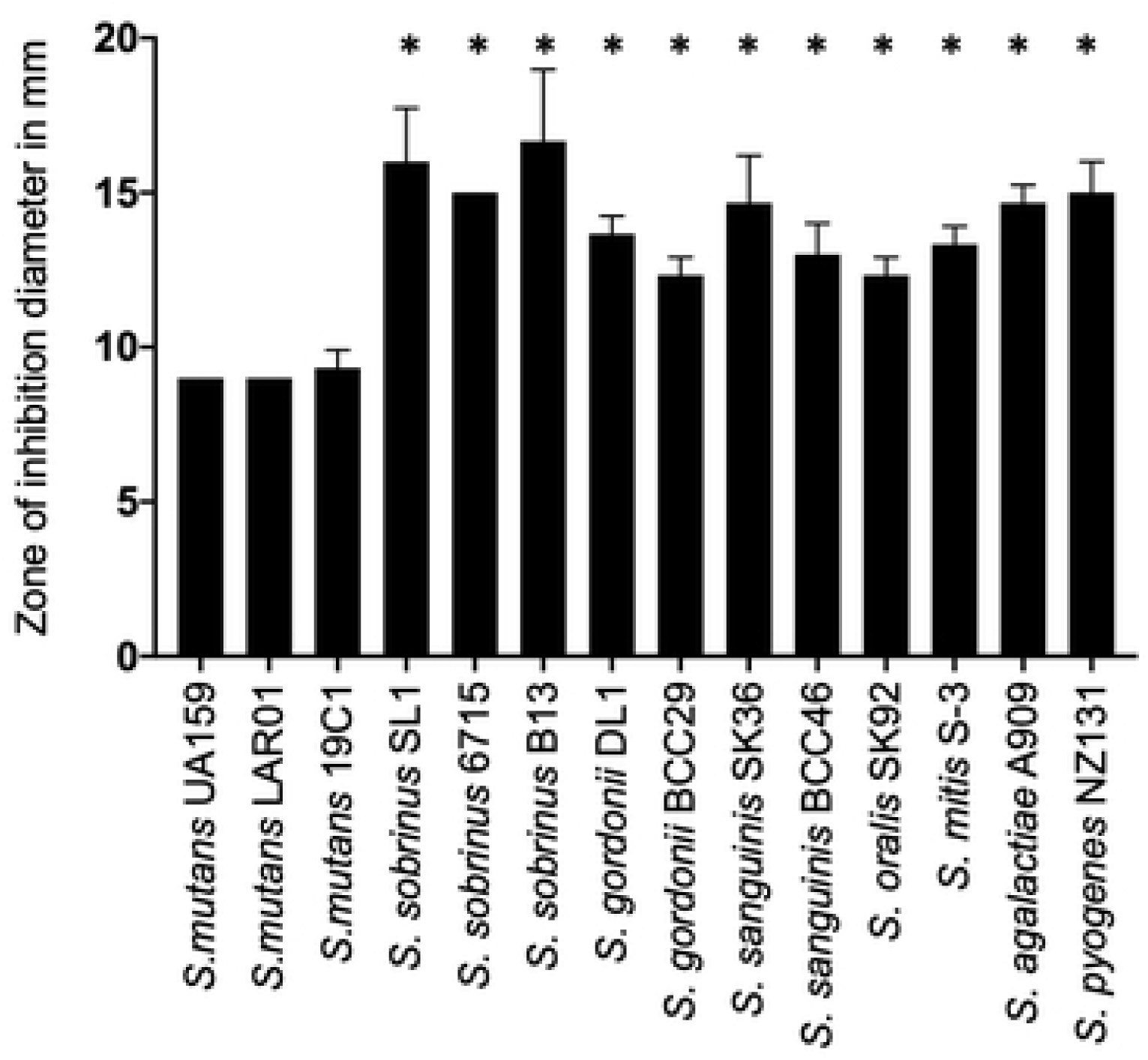
Disc diffusion assay indicating that *S. mutans* is inherently more tolerant to Zn than are other streptococci. Cultures were grown to mid-exponential phase, spread onto BHI plates, and topped with filter paper discs saturated with 1 mM ZnSO_4_. Growth inhibition zones (in mm) around the discs were measured after 24 h incubation at 37°C in 5% CO_2_. A *p* value of <0.05 was considered significant (*).

**Table 1.** ZnSO_4_ MIC values of different streptococcal strains.

| Species | Strain | ZnSO<sub>4</sub> MIC |
| --- | --- | --- |
| *S. mutans* | UA159 | 4 mM |
| *S. mutans* | LAR01 | 4 mM |
| *S. mutans* | 19C1 | 4 mM |
| *S. mutans* | 29SS | 4 mM |
| *S. sobrinus* | SL1 | 1 mM |
| *S. sobrinus* | 6715 | 1 mM |
| *S. sobrinus* | B13 | 1 mM |
| *S. gordonii* | DL1 | 0.5 mM |
| *S. gordonii* | K43BH3 | 0.5 mM |
| *S. sanguinis* | 18BH1 | 0.5 mM |
| *S. sanguinis* | SK36 | 0.5 mM |
| *S. oralis* | SK92 | 1 mM |
| *S. mitis* | S-3 | 1 mM |
| *S. intermedius* | ATCC27325 | 0.5 mM |
| *S. agalactiae* | A909 | 1 mM |
| *S. pyogenes* | NZ131 | 0.5 mM |
| *S. mutans* | UA159ΔzccE | 0.2 mM |
| *S. mutans* | UA159Δcop | 4 mM |
| *S. mutans* | UA159ΔzccEΔcop | 0.2 mM |

### Overview of changes to the *S. mutans* UA159 transcriptome following exposure to excess Zn

To obtain mechanistic insight into the attributes that promote the high Zn tolerance of *S. mutans*, we used RNA deep sequencing (RNA-Seq) to uncover the transcriptome of *S. mutans* UA159 following exposure to excess ZnSO_4_. To determine the most appropriate concentration of Zn for this experiment, the growth kinetics of exponentially grown cultures (OD_600_ ∼0.3) of *S. mutans* UA159 treated with increasing concentrations of ZnSO_4_ were monitored (**Fig S1**). As exposure of exponentially grown cultures to 4 mM ZnSO_4_ resulted in a marked growth defect, but not complete growth arrest as seen at 6 mM ZnSO_4_, this concentration was chosen for the transcriptome analysis. Applying a False Discovery Rate (FDR) of 0.05 and 2-fold cutoff, we found 96 genes upregulated and 62 genes downregulated after 15 minutes (T_15_) of exposure to 4 mM ZnSO_4_ when compared to the unstressed control (**Table S1**). After 90 minutes (T_90_), the number of differently expressed genes increased slightly from 158 at T_15_ to 199 genes, with 85 genes upregulated and 114 genes downregulated (**Table S1**). Among the genes differently expressed, 60 genes displayed the same expression trends (up or downregulated) at both time points whereas 9 genes showed opposing trends of regulation. To illustrate these results, differentially expressed genes were grouped according to Clusters of Orthologous Groups (COG) functional categories (**Fig 2**) (39).

**Fig 2.**
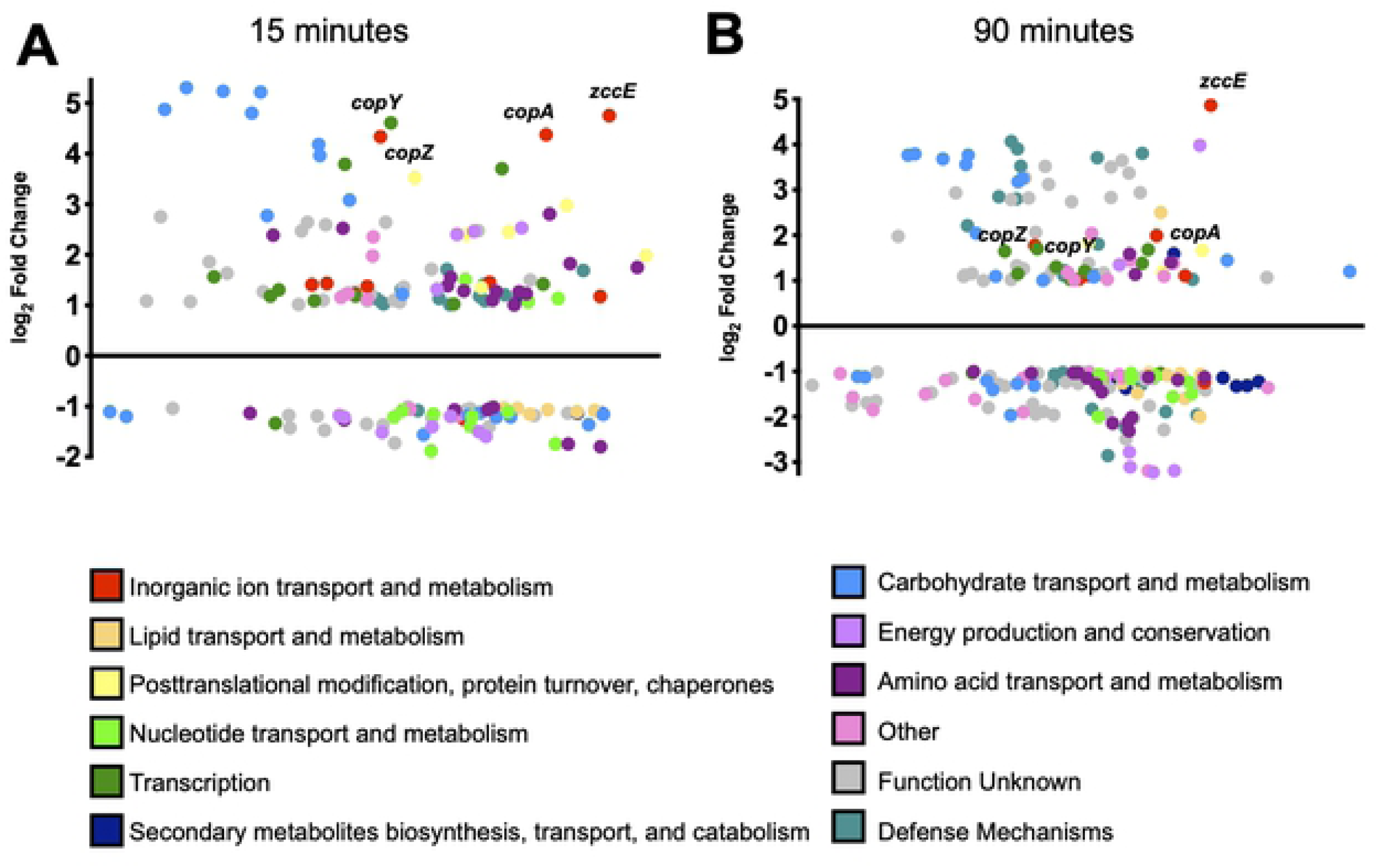
Summary of RNA-Seq analysis of *S. mutans* UA159 treated with 4 mM ZnSO_4_ for 15 (A) or 90 (B) minutes compared to untreated control. The y-axis indicates the log_2_ fold change in expression compared to the control, while the x-axis separates genes according to their average expression levels as compared to all other genes. The identities of selected genes of interest are indicated. The differentially expressed genes from both time points were grouped according to Clusters of Orthologous Groups (COG) functional categories (39).

One of the most highly induced (greater than 25-fold upregulation) genes at both time points was *smu2057c,* coding for a putative P_1B_-type ATPase transmembrane protein. Also, of interest, the entire *copYAZ* operon, of which *copA* encodes another P_1B_-type ATPase involved in Cu tolerance (40–42) were among the most upregulated genes at T_15_ (20- to 24-fold upregulated) remaining induced to a lesser extent at T_90_ (3- to 4-fold upregulated). Another group of genes highly upregulated at both time points encode for components of the lactose/galactose phosphoenolpyruvate transferase (PTS) and tagatose pathways (*lacXGEFDCBAR*, 4- to 40-fold upregulated). While genes associated with lactose uptake and utilization were strongly induced by high Zn stress, genes coding for the main PTS enzymes responsible for the uptake of sucrose (*scrA*), glucose (*manLMN*), and glucose disaccharide (*celCRB*) were downregulated at both time points. In addition, at T_15_, genes involved in the uptake or metabolism of cysteine (*tycABC*, *tcyEFGH*, *cysK*) and glutathione (*gst*, *gshT*, *gor*) were upregulated by 2- to 5-fold whereas genes encoding for mutacins IV (*nlmD*), V (*nlmC*) and VI (*nlmAB*) were induced by 8- to 16-fold at T_90_. Finally, several stress genes were induced at T_15_, including genes coding for heat shock proteins/molecular chaperone (*groES*-*EL* and *hrcA*-*grpE*-*dnaK*-*dnaJ* operons) and oxidative stress genes (*dpr*, *gor* and *tpx*). In contrast, the same *dpr* and *tpx* genes induced at T_15_ and additional genes classically associated with oxidative stress responses (*ahpCF*, *nox* and *sodA*) were downregulated at T_90_.

### *smu2057c* encodes a multi-metal translocating P-type ATPase unique to *S. mutans*

As *smu2057c* was one of the most strongly upregulated genes during high Zn stress and considering that bacterial P_1B_ -type ATPase proteins are implicated in metal ion transport (13–15, 43), we predicted that Smu2057c would function as a Zn efflux system. Compared to previously characterized P_1B_-type ATPases, the protein coded by *smu2057c* shares 33% amino acid identity with the *B. subtilis* CadA, which mediates tolerance to cadmium (Cd), Co and Zn (14, 44), and 36% identity to *E. coli* ZntA that mediates tolerance to Cd, lead (Pb) and Zn (13, 15) (**Fig 3**). Based on searches of public databases, *smu2057c* was exclusively found in *S. mutans* genomes, with exception of a single *Streptococcus troglodytae* genome encoding a hypothetical protein that was 94% identical to Smu2057c (**Table S2**).

**Fig 3.**
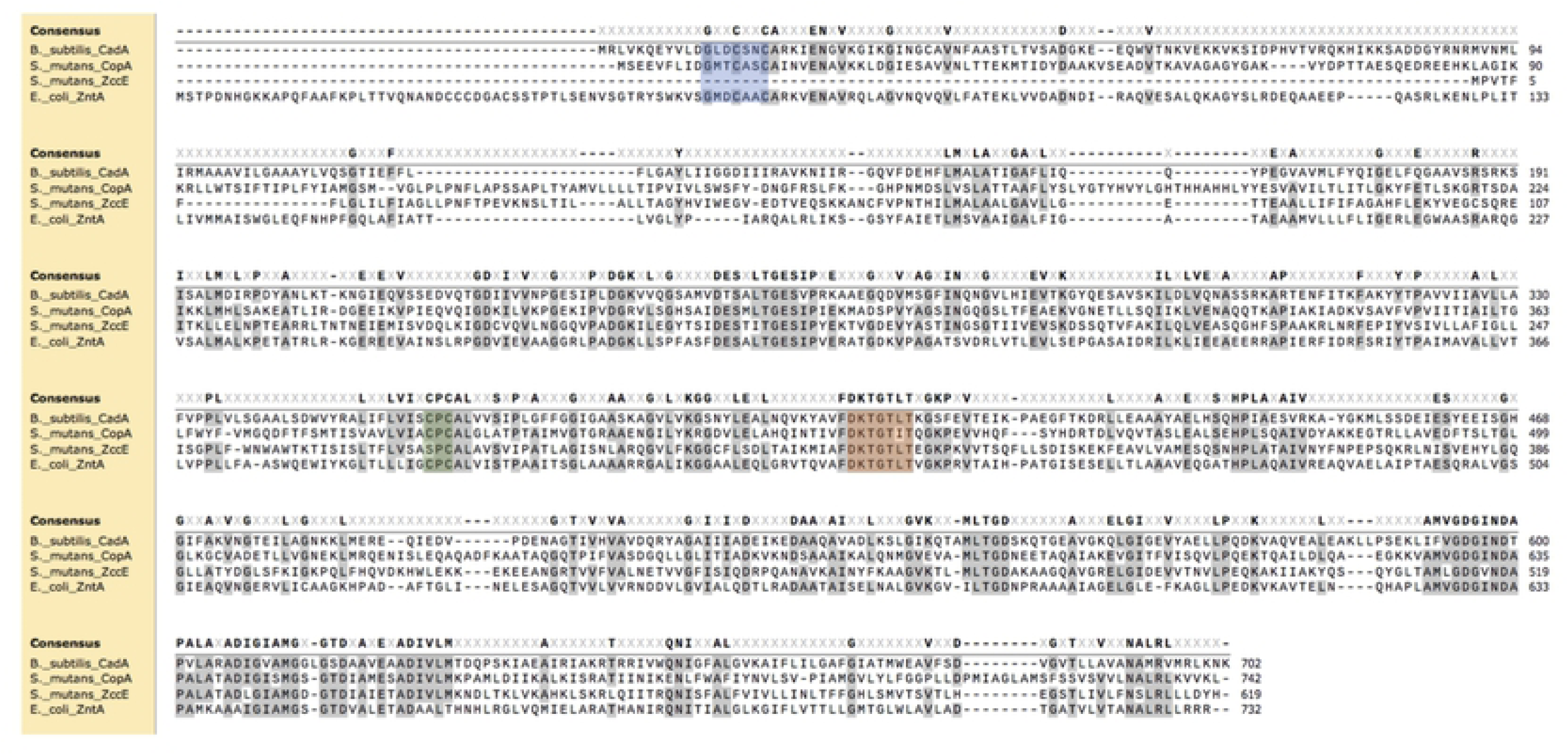
Amino acid sequence alignment of *S. mutans* P-type ATPases Smu2057c (ZccE) and CopA and P-type ATPases that confer Zn tolerance to *B. subtilis* (CadA) and *E. coli* (ZntA). Blue shade depicts the N-terminal metal binding motif that is absent in ZccE, green shade depicts the metal binding ‘CPC’ motif, and the orange shade indicates the conserved phosphorylation site for ATPase activity.

To determine if Smu2057c mediates Zn tolerance, we generated a *smu2057* deletion strain in the UA159 wild-type background and then tested the capacity of the mutant to grow in chemically-defined (FMC) or complex (BHI) media containing increasing concentrations of Zn. Because the experiments described below implicate Smu2057c in Cd, Co, Cu and Zn tolerance, we named *smu2057c* as *zccE*, for <u>z</u>inc, <u>c</u>admium, <u>c</u>obalt, and <u>c</u>opper <u>e</u>xporter. When compared to UA159, the Δ*zccE* strain displayed a much higher sensitivity to Zn as it was completely unable to grow in media supplemented with 1 mM ZnSO_4_ (**Fig 4A-C**). In agreement with the growth curves and plate titration assay, the Zn MIC of Δ*zccE* was 20 times lower than the MIC of the parent strain (**Table 1**). Genetic complementation of Δ*zccE* (Δ*zccE*^Comp^ strain) fully rescued the Zn sensitive phenotype (**Fig 4A-C**), while inactivation of *zccE* in the *S. mutans* OMZ175 background strain led to comparable and drastic reduction in Zn tolerance supporting that ZccE is the primary Zn tolerance determinant of *S. mutans* (**Fig S2**). Because P-type ATPases implicated in Zn tolerance have been shown to confer resistance to other metal ions, especially those of similar size and charge, we tested the capacity of Δ*zccE* to grow on BHI plates supplemented with Cd, Co, Cu, Fe, Mn and Ni. As shown in **Figure 4D**, ZccE confers resistance to Cd, Co and Cu but not to Fe, Mn or Ni.

**Fig 4.**
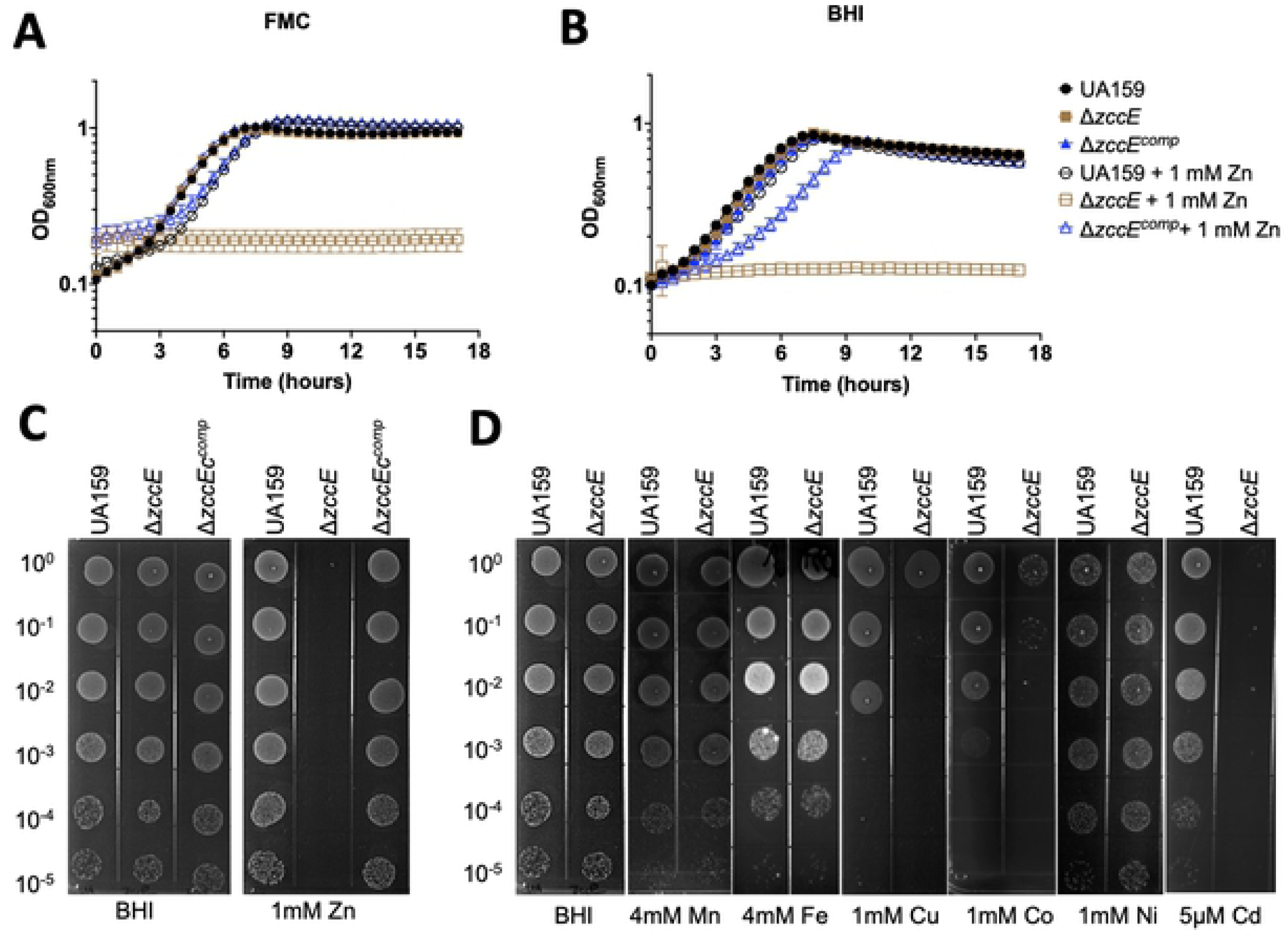
ZccE is a multi-metal exporter responsible for the high Zn tolerance of *S. mutans.* (A-B) Growth curves of *S. mutans* UA159, Δ*zccE* and Δ*zccE*^comp^ strains in FMC (A) or BHI (B) with or without 1 mM Zn supplementation. Data shown are average and standard deviation of 3 independent biological replicates. (C) Plate titration (spot test) of *S. mutans* UA159, Δ*zccE* and Δ*zccE*^comp^ strains on BHI agar with or without 1 mM Zn supplementation. (D) Plate titration of *S. mutans* UA159 and Δ*zccE* strains on BHI agar supplemented with 4 mM Mn, 4 mM Fe, 1 mM Cu, 1 mM Ni or 5 μM Cd. (C-D) Images were taken after 24 h incubation at 37°C in 5% CO_2_ and are representative of at least 3 independent experiments.

### ZccE and CopA work synergistically to protect *S. mutans* against Cu intoxication in oxidizing environments

In several bacteria, including all streptococci, Cu tolerance is mediated by the Cu- translocating P-type ATPase encoded by the *copA* gene, which is organized in an operon with *copY* (Cu-sensing transcriptional repressor) and *copZ* (Cu chaperone) (45–47). Because we found that ZccE also confers Cu tolerance and that the *copYAZ* operon was induced upon high Zn stress, we sought to determine to what extent ZccE contributes to Cu tolerance and to probe the possible association of CopA with Zn tolerance. To this end, we deleted the entire *copYAZ* operon in both the UA159 and Δ*zccE* background strains to generate the Δ*copYAZ* and Δ*zccE*Δ*copYAZ* strains. Next, we tested the capacity of each mutant strain to grow in media containing high concentrations of CuSO_4_ or ZnSO_4_. Despite strong upregulation of the *copYAZ* operon during the initial 15 minutes of high Zn stress (**Table S1, Fig 2**), inactivation of *copYAZ* alone did not affect Zn sensitivity (**Fig 5A**). While the Δ*zccE*Δ*copYAZ* strain phenocopied Δ*zccE* on plates containing 0.1 or 1 mM ZnSO_4_, the double mutant was slightly more sensitive than the Δ*zccE* single mutant on plates containing 0.05 mM ZnSO_4_ (**Fig 5A**). Collectively, these results indicate that CopA plays a minor role in Zn tolerance that becomes dispensable when ZccE is present.

**Fig 5.**
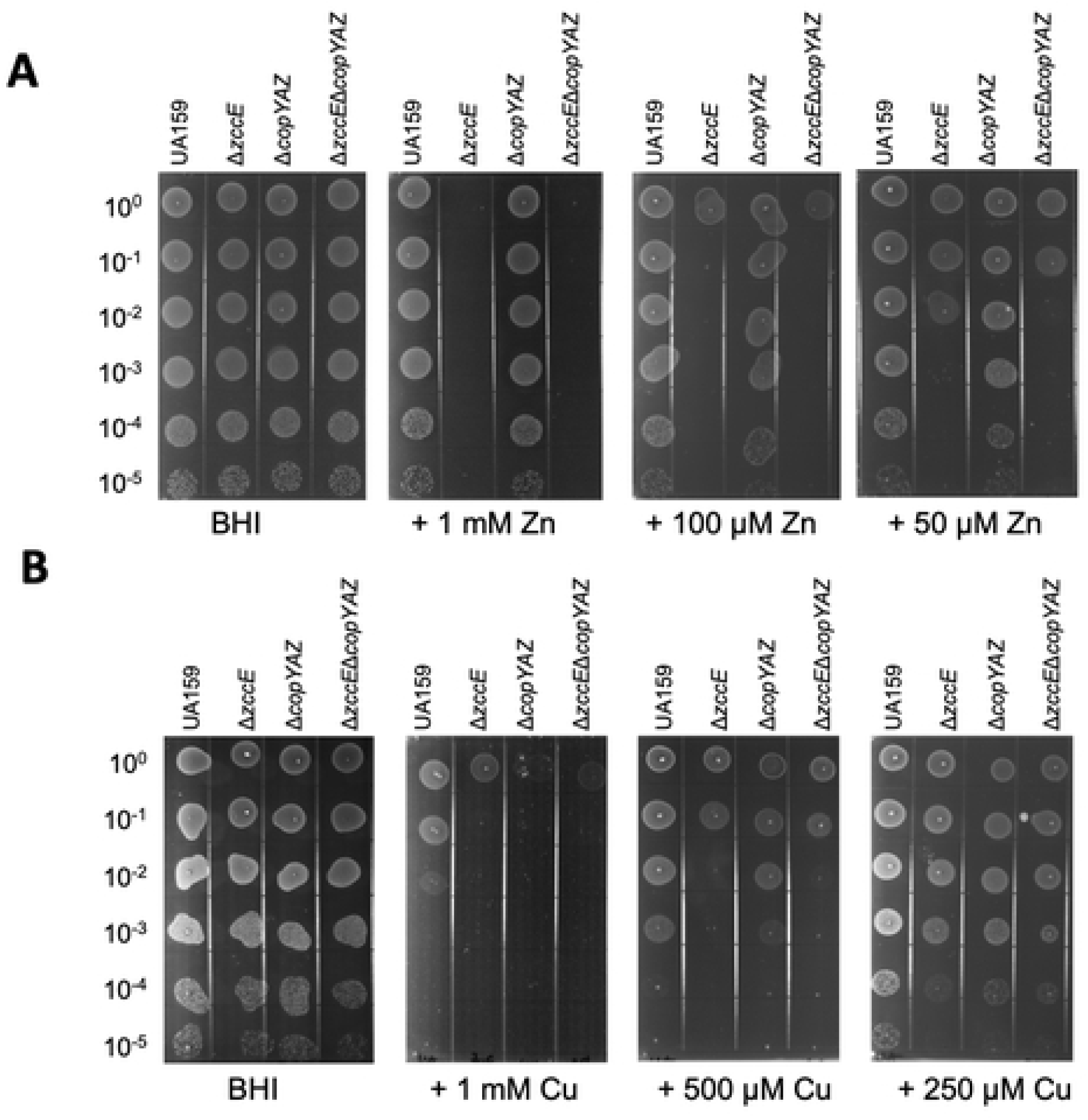
CopA plays a minor role in Zn tolerance while ZccE and CopA work interchangeably to protect against Cu intoxication in oxidizing environments. (A-B) Plate titration (spot test) of *S. mutans* UA159, Δ*zccE*, Δ*copYAZ*, and Δ*zccE*Δ*copYAZ* strains on BHI agar with varying concentrations of Zn (A) or Cu (B). Images were taken after 24 h incubation at 37°C in 5% CO_2_ and are representative of at least 3 independent experiments.

As shown by others (40, 45), inactivation of the *copYAZ* operon drastically heightened Cu sensitivity albeit the Δ*zccE* strain was slightly more sensitive than Δ*copYAZ* on plates containing 500 μM CuSO_4_. Unexpectedly, inactivation of both *copYAZ* and *zccE* did not appear to further enhance Cu sensitivity when compared to the single mutant strains. Though Cu should primarily be found as Cu^+^ in the reducing bacterial cytoplasm (48), it is capable of alternating between Cu^+^ or Cu^2+^. Thus, we wondered if ZccE could mobilize both oxidized and reduced Cu and, if so, whether there was a preference for one redox state over another. Of note, previous work has shown that the Pneumococcal Cop operon can efficiently remove Cu^+^ and Cu^2+^ from the bacterial cytoplasm into the extracellular milieu (49). In light of this redox cycling, we reevaluated the Cu tolerance of Δ*zccE* and Δ*copYAZ* single and double mutants under anaerobiosis such that all Cu ions available should be on the reduced Cu^+^ form. In agreement with the knowledge that Cu^+^ is more toxic than Cu^2+^, supplementation of agar plates with 1 mM CuSO_4_ caused complete growth inhibition of all strains (data not shown), such that experiments using an anaerobic environment were conducted with plates containing at most 0.1 mM CuSO_4_. Interestingly, we found that Cu tolerance under anaerobiosis was solely mediated by CopA, suggesting that ZccE cannot mobilize Cu^+^ (**Fig S3**). Taken together, these results reveal that CopA and ZccE interchangeably mediate Cu protection in aerobic environments. However, CopA is likely to have greater biological relevance to Cu tolerance than ZccE due to its unique capacity to export Cu^+^.

### ZccE is positively regulated by a MerR-type regulator

Sequence analysis revealed that *zccE* is located immediately upstream but transcribed in the opposite orientation of *smu2058*, an uncharacterized MerR-type transcriptional regulator (**Fig 6A**). To explore a possible role of Smu2058 in *zccE* regulation, the *smu2058* coding sequence was replaced by a Spec^R^ cassette generating the Δ*zccR* strain (for *<u>z</u>ccE* regulator). Using the same growth conditions as were used in the RNA-Seq analysis described above, quantitative real time PCR (qRT-PCR) analysis revealed that *zccE* transcription is almost entirely dependent on ZccR (**Fig 6B**). In agreement with this observation, the Δ*zccR* strain phenocopied the Δ*zccE* strain and was unable to grow in broth (**Fig S4**), or on agar plates (**Fig 6C**) supplemented with ZnSO_4_. Genetic complementation of Δ*zccR* (Δ*zccR*^Comp^ strain) reversed the heightened Zn sensitivity indicating that the phenotype was not due to polar effects on *zccE* or secondary mutations (**Fig S4 and Fig 6C**).

**Fig 6.**
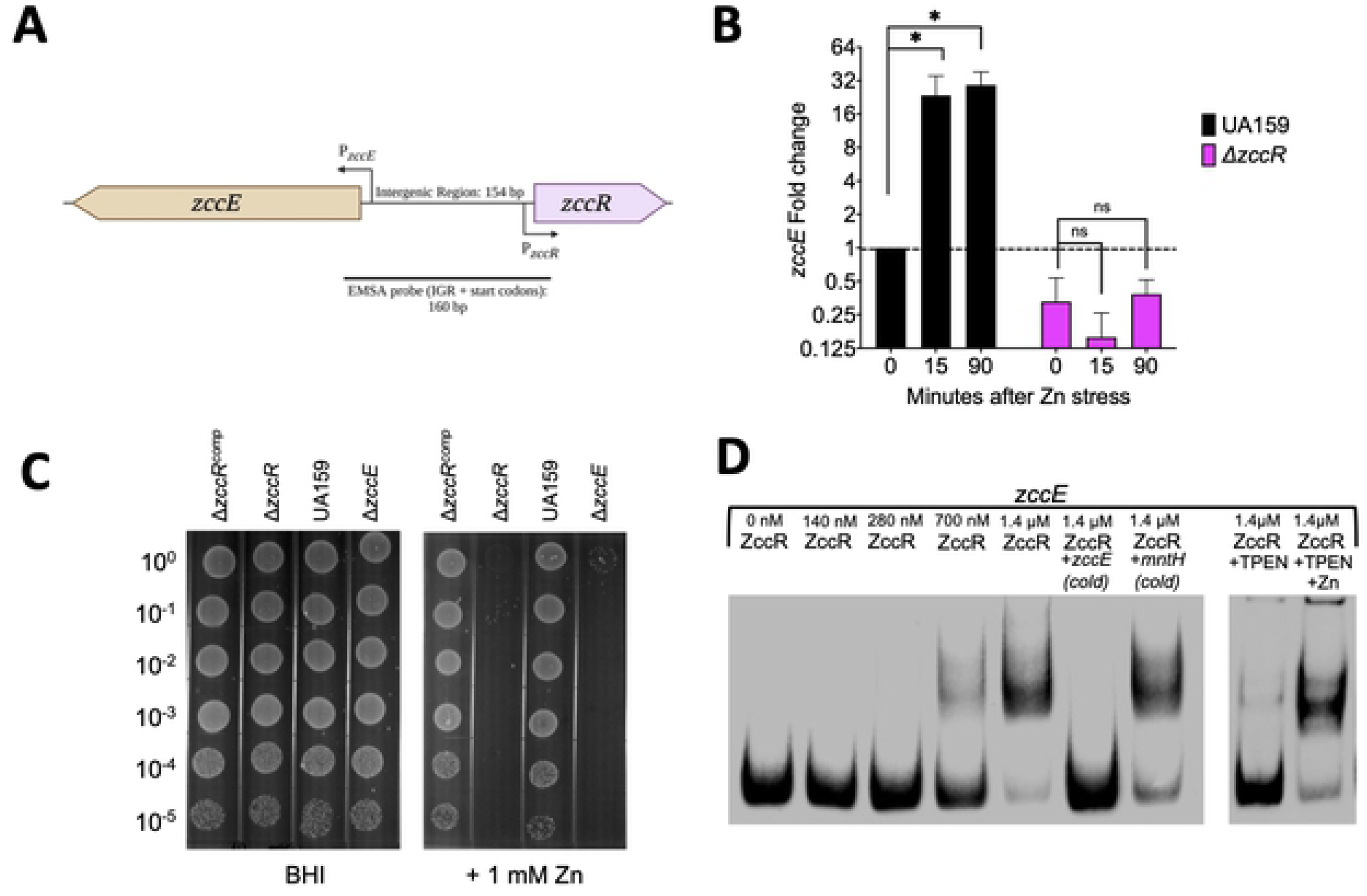
The MerR-type transcription factor ZccR is a positive regulator of *zccE*. (A) Genetic organization of *zccE* (*smu2057c*) and *zccR* (*smu2058*) in the *S. mutans* UA159 chromosome (Created with BioRender.com). (B) qRT-PCR analysis of *zccE* mRNA expression in the Δ*zccR* strain relative to the parent UA159 strain before and after exposure to 4 mM Zn. Data represent means and standard deviations of results from 3 independent experiments. One-way ANOVA was used to determine significance (*, *p* <0.001; ns, not significant). (C) Plate titration (spot test) of UA159, Δ*zccE*, Δ*zccR* and Δ*zccR^comp^* strains on BHI with or without 1 mM Zn supplementation. Images were taken after 24 h incubation at 37°C in 5% CO_2_ and are representative of at least 3 independent experiments. (D) EMSA showing direct interaction of ZccR with the 160-bp *zccE*- *zccR* intergenic region (IGR). To determine ZccR binding specificity, addition of 100X (molar excess) of either non-biotin labeled (cold) *zccE*-*zccR* IGR or non-specific competitor DNA (*mntH* promoter region) was added to the reaction. To explore the role of Zn on ZccR binding activity, 3 mM TPEN (Zn specific chelator) was added to the reaction alone or in the presence of 2.5 mM ZnSO_4_.

To determine if ZccR regulation of *zccE* transcription is direct or indirect, we performed electron mobility shift assays (EMSAs) using recombinant ZccR purified from *E. coli* and a DNA probe encompassing the entire *zccE*-*zccR* intergenic region (**Fig 6A**). The EMSA results reveal that addition of sufficient ZccR resulted in a shift of the protein: DNA complex, indicating that ZccR binds specifically to the *zccE*-*zccR* intergenic region (**Fig 6D**). Formation of the ZccR:DNA complex was inhibited by the Zn-specific chelator TPEN (*N*,*N*,*N′*,*N′*-tetrakis(2-pyridinylmethyl)- 1,2-ethanediamine) and competitively reverted by the excess Zn supplementation (**Fig 6D**). Thus, we conclude that ZccR positively and directly regulates *zccE* expression, likely by sensing and responding to intracellular Zn pools.

### Inactivation of *zccE* or *zccR* drastically alters Zn:Mn ratios in high Zn conditions

Next, we used inductively coupled plasma mass spectrometry (ICP-MS) to quantify the intracellular Mn and Zn pools in the UA159 (parent strain), Δ*zccE* and Δ*zccR* strains exposed to high Zn stress. Briefly, cultures were grown in BHI broth (∼ 10 µM Zn,∼ 0.5 µM Mn (50)) to mid-log phase and treated with a ZnSO_4_ solution to reach a final concentration of 1 mM of Zn. For ICP-MS analysis, samples were obtained immediately before (control) and 90 minutes after addition of Zn. Despite the high concentration of Zn added to the culture, parent and complemented mutant strains maintained the same intracellular amounts of Zn before and after the Zn challenge (**Fig 7A**). While intracellular levels of Zn in the Δ*zccE* and Δ*zccR* strains before addition of Zn did not differ from parent and complemented mutants, intracellular Zn pools rose by ∼10-fold in the mutants after Zn challenge (**Fig 7A**). Moreover, accumulation of Zn in the Δ*zccE* and Δ*zccR* strains inversely correlated with a substantial decrease (∼10-fold) in intracellular Mn pools (**Fig 7B**).

**Fig 7.**
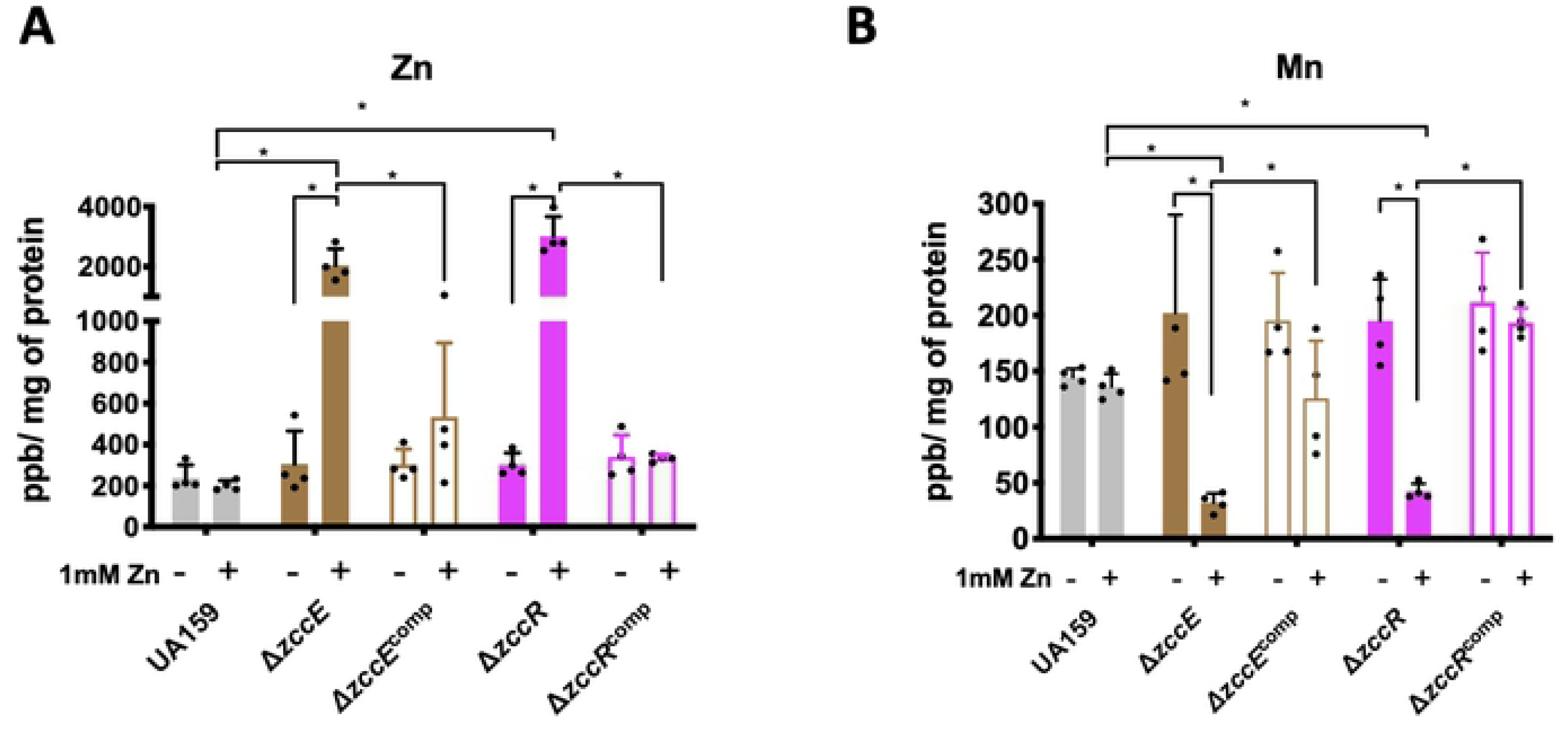
ICP-MS quantifications of intracellular Zn and Mn pools in UA159, Δ*zccE*, Δ*zccR* and respective complemented strains. Strains were grown in BHI to mid-exponential phase at which point the experimental groups were exposed to 1 mM Zn for 90 minutes. The graphs indicate intracellular parts per billion Zn (A) or Mn (B) normalized by milligram of total protein. Data represent average and standard deviation of values from four independent biological replicates. Two-way ANOVA was used to determine significance between metal content of either the same strain before and after Zn exposure or among different strains after Zn exposure. A *p* value of <0.05 was considered significant (*).

### Disruption of Zn homeostasis diminishes oxidative stress tolerance

As both Zn and Mn are linked to oxidative stress responses (50–52), we sought to determine if simultaneous perturbations of Zn and Mn homeostasis (and therefore of Zn:Mn ratios) in Δ*zccE* affected the ability of this strain to cope with peroxide stress. In disc diffusion assays, both UA159 and Δ*zccE* strains displayed similar sensitivity to H_2_O_2_ when grown in BHI alone(∼ 10 µM Zn) while supplementation of sub-inhibitory concentrations of Zn (50 to 100 μM ZnSO_4_) resulted in significantly greater sensitivity of Δ*zccE* when exposed to H_2_O_2_ (**Fig 8A**). Next, we tested the ability of *S. gordonii* DL1, a peroxigenic oral commensal, to inhibit the growth of the *S. mutans* parent and Δ*zccE* strains using an antagonism plate assay. Like the disc diffusion assay, both strains showed similar sensitivities to *S. gordonii* when grown on BHI plates, while supplementation of the media with as little as 50 μM ZnSO_4_ virtually abolished growth of Δ*zccE* (**Fig 8B**). To confirm that the growth defect was due to sensitivity to the H_2_O_2_ produced by *S. gordonii*, a control assay was performed in which catalase was added atop the *S. gordonii* prior to spotting of the *S. mutans* cultures. When the H_2_O_2_ was neutralized by catalase, no growth sensitivity by the *S. mutans* strains was observed (**Fig 8B**).

**Fig 8.**
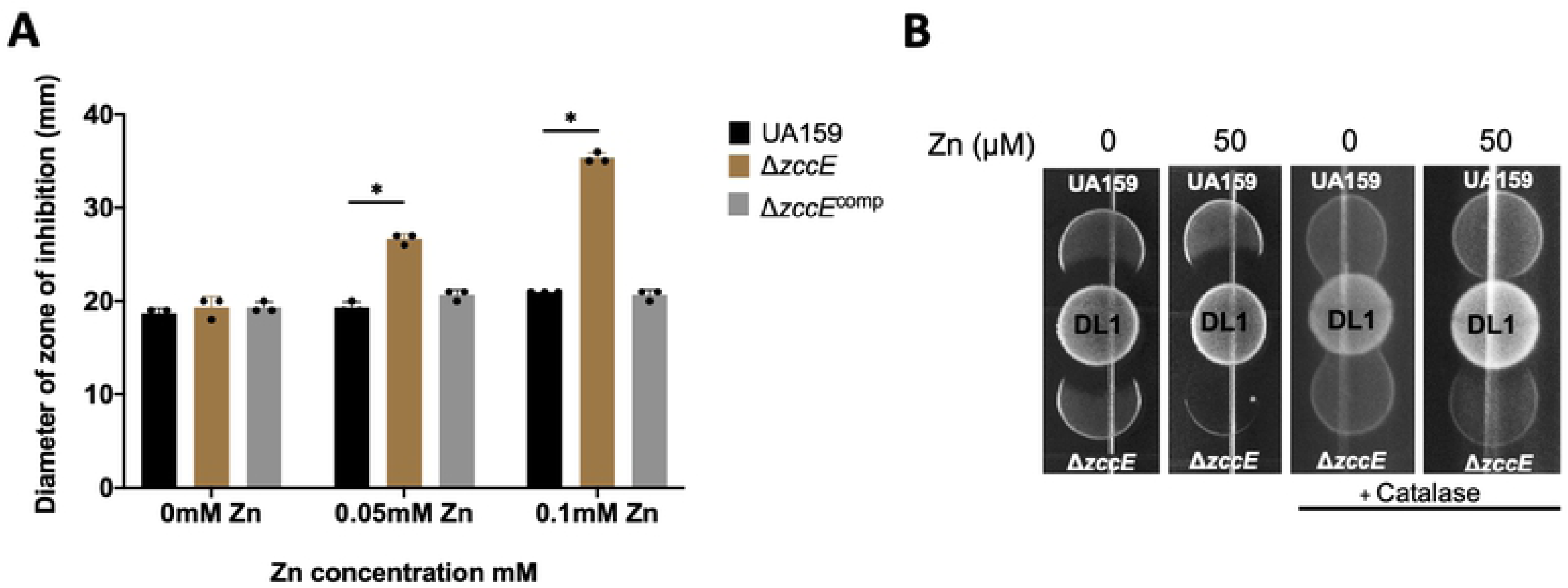
Disruption of Zn homeostasis increases H_2_O_2_ sensitivity in the *zccE* strain. (A) Growth inhibition zones for *S. mutans* UA159, Δ*zccE*, and Δ*zccE*^Comp^ strains grown on BHI agar with or without Zn supplementation and exposed to filter paper discs saturated with 0.25% H_2_O_2_. Two- way ANOVA was used to determine significance (*, *p* <0.05). (B) Growth inhibition zones of *S. mutans* UA159 and Δ*zccE* strains by peroxigenic *S. gordonii* DL-1 spotted on BHI agar with or without 50 μM Zn supplementation. The *S. mutans*-*S. gordonii* competition assay was repeated with catalase overlaid onto the *S. gordonii* spot to inactivate H_2_O_2_. Images are representative of results from three independent experiments.

### Topical Zn treatment impairs oral colonization efficiency of the **Δ*zccE* strain**

To determine the impact of Zn homeostasis in an *in-vivo* model, we assessed the ability of the parent and Δ*zccE* strains to colonize the oral cavity of rats fed a cariogenic diet, while simultaneously testing whether topical Zn treatment could inhibit bacterial colonization (**Fig 9**). To simulate oral hygiene such as a mouthwash, rats that had been orally infected with *S. mutans* UA159 or *ΔzccE* were treated twice daily with an oral application of a ZnSO_4_ solution. When comparing the control treatment groups (saline), similar numbers of colony-forming units (CFU) were recovered from animals infected with either UA159 or Δ*zccE*, indicating that the multi-metal tolerance conferred by ZccE was dispensable for oral colonization in the absence of excess Zn. However, daily treatment with 60 mM or 150 mM ZnSO_4_ solutions, a concentration range found in commercially available oral healthcare products, resulted in significant reduction (∼1-log) of the CFU recovered from rats that had been infected with the Δ*zccE* strain. These same Zn treatments showed only modest, and not statistically significant, inhibitory effects against the parent strain. Importantly, Zn treatment did not affect the abundance of total flora recovered from the *S. mutans*- infected animals, suggesting that the inhibitory effect was specific to the Δ*zccE* strain. Taken together, these results provide the proof of concept that ZccE is a suitable target for future development of an antimicrobial therapy specific to treat *S. mutans* infections.

**Fig 9.**
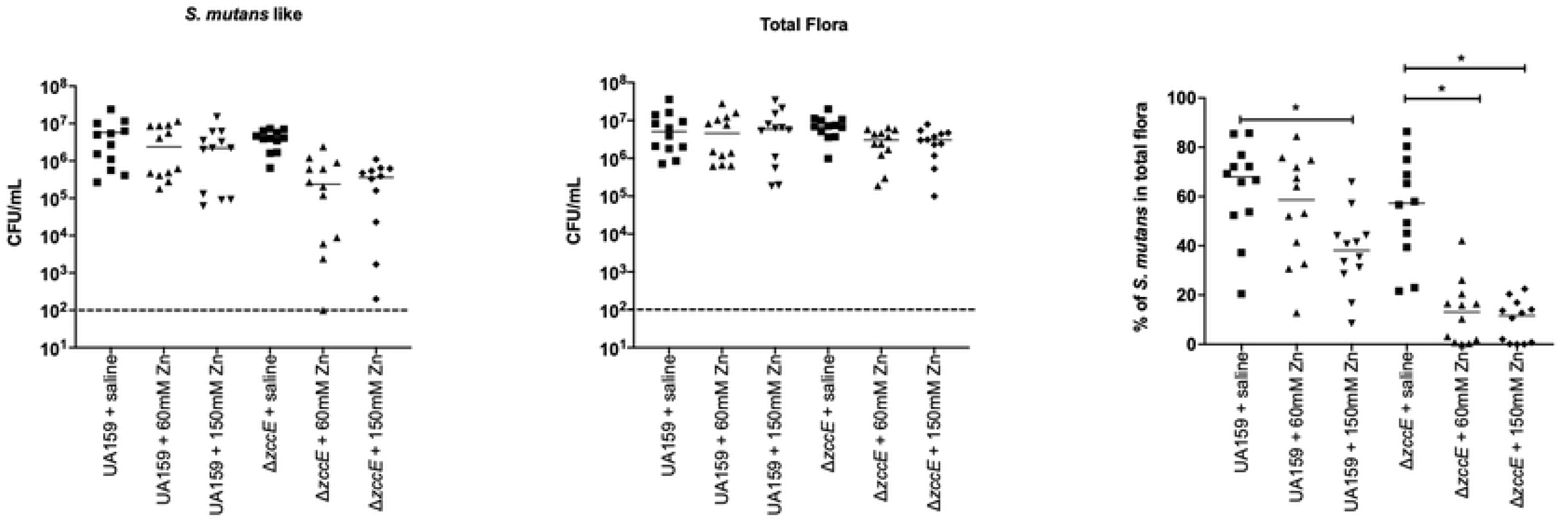
Colonization efficiency of *S. mutans* UA159 or Δ*zccE* on the teeth of rats after topical treatment with Zn. (A-B) Bacterial CFU recovered from rat jaws by plating on (A) MS agar for *S. mutans* and (B) blood agar for total flora. (C) Percentage of *S. mutans* colonies were calculated by dividing the values shown in (B) by those shown in (A). The dashed line indicates the limit of detection. One-way ANOVA was used to determine significance. A *p* value of <0.05 was considered significant (*).

## Discussion

The second most abundant trace metal in the human host, Zn is naturally found in saliva and dental plaque at levels that, in theory, should not be limiting or toxic to bacteria. Yet, our recent characterization of the AdcABC Zn import system suggested that Zn might be a growth-limiting factor in dental biofilms, as inactivation of the *adcABC* metal transport system greatly reduced *S. mutans* colonization in a rat oral infection model (38). Whether this apparent Zn restriction was caused by host-mediated Zn sequestration mechanisms, increased competition for environmental Zn with microbial commensals, or the combination of both remains to be determined. In addition, Zn is required for host immune function and shows antimicrobial properties at elevated concentrations, such that it is harnessed by the host to intoxicate invading pathogens within phagocytic cells and, for centuries, has been used as a therapeutic agent to treat a variety of human conditions due to its antimicrobial and anti-inflammatory properties. In oral health, Zn is incorporated into oral healthcare products to treat gingivitis and to control halitosis (27, 28) though its usefulness to caries treatment or prevention has proven to be controversial (27, 33, 53, 54). In this study, we used transcriptomic and molecular genetic approaches to uncover the mechanisms that allow *S. mutans* to overcome Zn toxicity. Through RNA-Seq analysis, we identified a previously uncharacterized P-type ATPase transporter, encoded by *smu2057c* (*zccE*), as involved in Zn tolerance. Importantly, we discovered that *zccE* is part of the core genome and unique to *S. mutans*. By comparing Zn tolerance levels of several streptococcal species that do not encode ZccE with those of the wild-type and Δ*zccE* strains of *S. mutans*, it became clear that ZccE was directly responsible for the inherently high Zn tolerance of *S. mutans*. Like other bacterial metal-translocating P-type ATPases that have the capacity to translocate multiple metals, loss of *zccE* also led to increased sensitivity to Cd, Co and Cu. In addition, we identified *zccR* (*smu2058*), a member of the MerR family of transcriptional factors, as the major transcription factor controlling *zccE* expression in a Zn-dependent manner.

In addition to identification of *zccE*, the RNA-Seq analysis provided additional clues on how *S. mutans* might overcome Zn intoxication. For example, transcriptional changes in the expression of sugar transport and sugar metabolism genes suggest that *S. mutans* activates the tagatose-6-phosphate pathway to bypass the initial steps of the glycolytic pathway. These steps have been shown to be inhibited by millimolar concentrations of Zn in the closely related species *S. pyogenes* (Ong, Walker, & McEwan, 2015). Specifically, our transcriptome results revealed that elevated Zn concentrations repressed transcription of PTS genes involved in glucose and glucose disaccharide metabolism whereas genes of the lactose PTS and the tagatose pathway were strongly induced (**Table S1 and Fig 2**). A well-known mechanism utilized by bacteria to overcome Zn toxicity is based on the accumulation of Zn-buffering molecules including cysteine-containing molecules such as glutathione and free cysteine (8, 55). Our RNA-Seq analysis also revealed that *S. mutans* upregulates expression of genes involved in glutathione transport (*tcyABC*) and regeneration (*gst*), cystine transport (*tcyDEFG*) and metabolism (*cysK*) in response to high Zn stress. Future studies are necessary to test the hypothesis that *S. mutans* activates lactose metabolism to bypass a possible metabolic bottleneck created by Zn-mediated inhibition of the initial steps of glycolysis, and to whether glutathione and cysteine are indeed used as Zn- buffering systems to protect *S. mutans* against Zn intoxication.

Despite the multiple strategies bacteria utilize to mitigate Zn poisoning, it is the capacity to efficiently pump Zn out of their cytosol that seems to primarily define the levels of Zn tolerance. Until now, the only Zn efflux system known in streptococci was encoded by *czcD*, a CDF-type efflux system that has been characterized in major human pathogens including *S. pneumoniae*, *S. pyogenes*, and *S. agalactiae* (9, 18, 19, 22). However, the genome of *S. mutans* does not encode a *czcD* homologue; the only *S. mutans* gene coding for a CDF-type transporter is *smu1176* (*mntE*), which was recently shown to alleviate Mn toxicity (56). Of interest, several bacteria, including the Gram-positive paradigm *B. subtilis*, have been shown to utilize both a CDF-type transporter and a P-type ATPase transporter to maintain Zn homeostasis during high Zn stress (12, 14).

P-type ATPases are a large family of transmembrane proteins that use the energy generated by ATP hydrolysis to transport cations and other substrates across membranes. P_1B_- type ATPases are primarily involved in metal ion transport and are comprised of six to eight transmembrane helices that form the ionic channel across the cell membrane with two soluble cytoplasmic domains. In these metal translocating P-type ATPases, one of the eight helices bears a tripeptide ‘CPC’ metal-binding signature motif with a conserved center proline flanked by either cysteine/serine or histidine at either side [(C**/**S/T)P(C**/**H/P)] (57, 58). While the proline residue is essential, the amino acids surrounding the proline have been shown to define metal specificity of these transmembrane proteins. For example, the canonical *E. coli* ZntA has a fully conserved ‘CPC’ motif that confers high selectivity to Pb followed by Zn and Cd, and low selectivity to Co, Cu and Ni – substitution of one of the surrounding cysteines by histidine or serine resulted in loss of binding to Pb but not to Zn or Cd (59). Based on previous studies, P_1B_-type ATPases with ‘CPC’ motifs are selective for silver (Ag), Cd, Cu, Pb, and Zn whereas proteins with SPC or CPH motifs are associated with Co and Cu export (59). In addition to the ‘CPC’ motif, metal specificity is determined by the presence of GxxCxxC motif at the N-terminal end and the presence of a preceding aspartate residue (59). Although ZccE possesses an ‘SPC’ motif, the N-terminal GxxCxxC motif is absent in ZccE such that the coordination chemistry and key residues of ZccE that determine metal specificity or promiscuity remain to be fully determined.

One of the best described mechanisms by which Zn exerts its toxicity in bacteria is by impairing Mn homeostasis (18, 60). In *S. pneumoniae*, Zn binds to the Mn transporter PsaABC when present at high concentrations, competitively affecting Mn uptake and rendering *S. pneumoniae* hypersensitive to oxidative stress and immune cell killing (61). Of note, Zn outcompetes Mn for binding to PsaABC without being internalized (61). Follow up studies with *S. pneumoniae* and *S. agalactiae* indicated that a drastic alteration in Zn:Mn ratios, rather than Zn concentration itself, was the main cause for Zn toxicity; Mn supplementation alone restored the equilibrium of Zn:Mn ratios and alleviated the growth defect phenotypes associated with high Zn stress (22, 60, 61). Here, we found that the ∼10-fold increase in intracellular Zn levels in Δ*zccE* was accompanied by a similar decrease in intracellular Mn pools (**Fig 7**), such that Zn:Mn ratios changed from ∼1:1 before addition of Zn to ∼100:1 after the challenge. While Zn has been shown to compete with Mn for binding to the DtxR family transcriptional repressor SloR (62), our RNA-Seq analysis indicates that the decrease in total Mn pools was not caused by altered transcription of the SloR-regulated genes responsible for either Mn import (*sloABC* and *mntH*) or export (*mntE*) (50, 56). Mn plays multiple and important roles in oxidative stress tolerance, so this notable drop in intracellular Mn pools provides one possible explanation, albeit likely not the only one, for the hypersensitivity of Δ*zccE* to H_2_O_2_ when simultaneously exposed to sub-inhibitory concentrations of Zn (**Fig 8**). Of interest, the parent strain was capable of maintaining steady levels of both Zn and Mn (and therefore balanced Zn:Mn ratios) when challenged with 1 mM Zn, suggesting that intracellular and not extracellular Zn levels are affecting Mn homeostasis. To further explore the association of Mn with Zn toxicity effects, we asked if Mn supplementation could rescue phenotypes of the *ΔzccE* strain when exposed to excess Zn. When *ΔzccE* was exposed to high Zn stress, the addition of Mn offered only minimal growth restoration (**Fig S5A**). However, the restorative effect of Mn supplementation was clear in the context of tolerance of Δ*zccE* to H_2_O_2_ (**Fig S5B**). Taken together, these results support previous investigations that demonstrate that Zn and Mn homeostasis are intertwined and that this relationship is critical to bacterial pathophysiology. For this reason, mechanistic studies to understand how intracellular Zn perturbs Mn flux in *S. mutans* warrants further investigation.

To explore the potential of ZccE as an antimicrobial target, we performed an oral colonization study that included topical application of a ZnSO_4_.treatment to animals that had been infected with S. mutans UA159 or Δ*zccE*. As anticipated, the Δ*zccE* strain was susceptible to Zn treatment whereas the parent strain was not (**Fig 9**). These results suggest that targeted inhibition of ZccE expression or activity can be combined with Zn to develop an anti-caries therapy by specifically killing *S. mutans*. As a proof-of-principle, we tested the inhibitory effects of Zn and Na-orthovanadate, a P-type ATPase inhibitor (63), against *S. mutans* UA159. While orthovanadate is expected to inhibit the activity of other P-type ATPases (the *S. mutans* genome encodes four P-type ATPases), we observed a synergistic inhibitory effect of Zn and orthovanadate as evidenced by the MIC values shown in **Fig S6**.

To summarize, this report sheds new light onto the molecular mechanisms utilized by *S. mutans* to overcome Zn toxicity. We discovered that *S. mutans* is inherently more tolerant to the toxic effects of Zn than other streptococci, including several non-cariogenic oral streptococci that are associated with oral health. We also identified the major effectors of high Zn tolerance in *S. mutans*, the metal translocating P_1B_-type ATPase ZccE and the Zn-responsive transcriptional factor ZccR. Because ZccE is unique to *S. mutans*, we propose that small molecules that specifically inhibit ZccE activity (or expression) can be combined with a Zn source to kill *S. mutans*. Thus, the combination of a ZccE-specific inhibitor with Zn can be exploited as a new antimicrobial therapy to prevent dental caries or to treat systemic *S. mutans* infections.

## Materials and methods

### Bacterial strains and growth conditions

The strains used in this study are listed in **Table 2**. All strains were routinely grown in brain heart infusion (BHI) at 37°C in a 5% CO_2_ atmosphere. When appropriate, antibiotics were added to cultures at the following concentrations: spectinomycin (1 mg ml^-1^), kanamycin (1 mg ml^-1^), erythromycin (10 μg ml^-1^). BHI and the chemically-defined medium FMC (64) were used to generate growth curves using an automated growth reader (Bioscreen C; Oy Growth Curves Ab, Ltd.). Briefly, overnight cultures were sub-cultured 1:20 in fresh media, grown to mid-exponential phase (OD_600_ of 0.4), and diluted 1:50 into the appropriate medium with or without metal supplementation in the wells of a microtiter plate. An overlay of sterile mineral oil was added to each well to minimize generation of reactive oxygen species. For RNA-Seq and qPCR analyses, bacterial cultures were grown in BHI to mid-log phase (OD_600_ of 0.4, untreated control) and then exposed to a final Zn concentration of 4 mM (using a concentrated ZnSO_4_ solution) for 15 or 90 minutes.

**Table 2.**
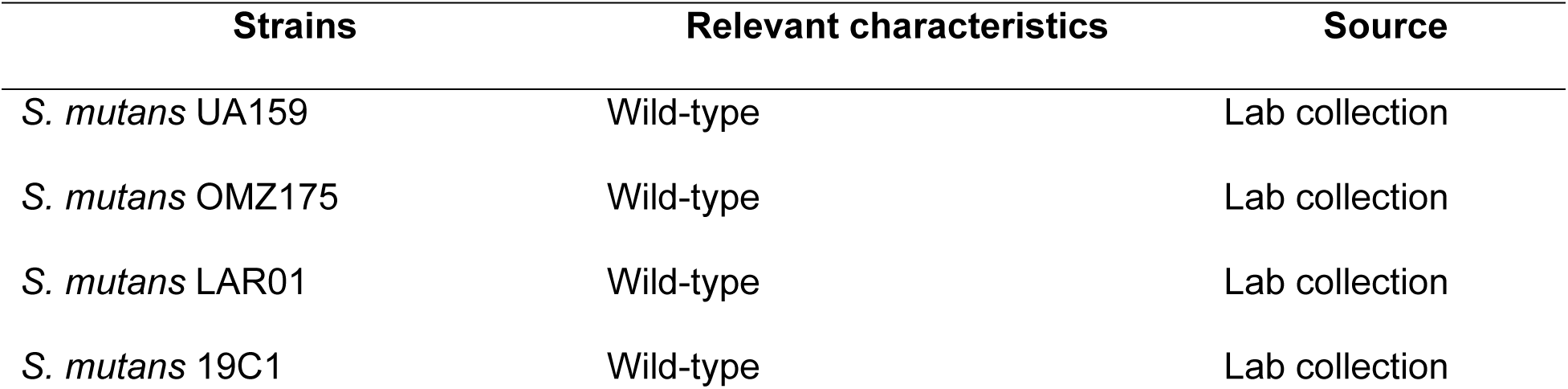
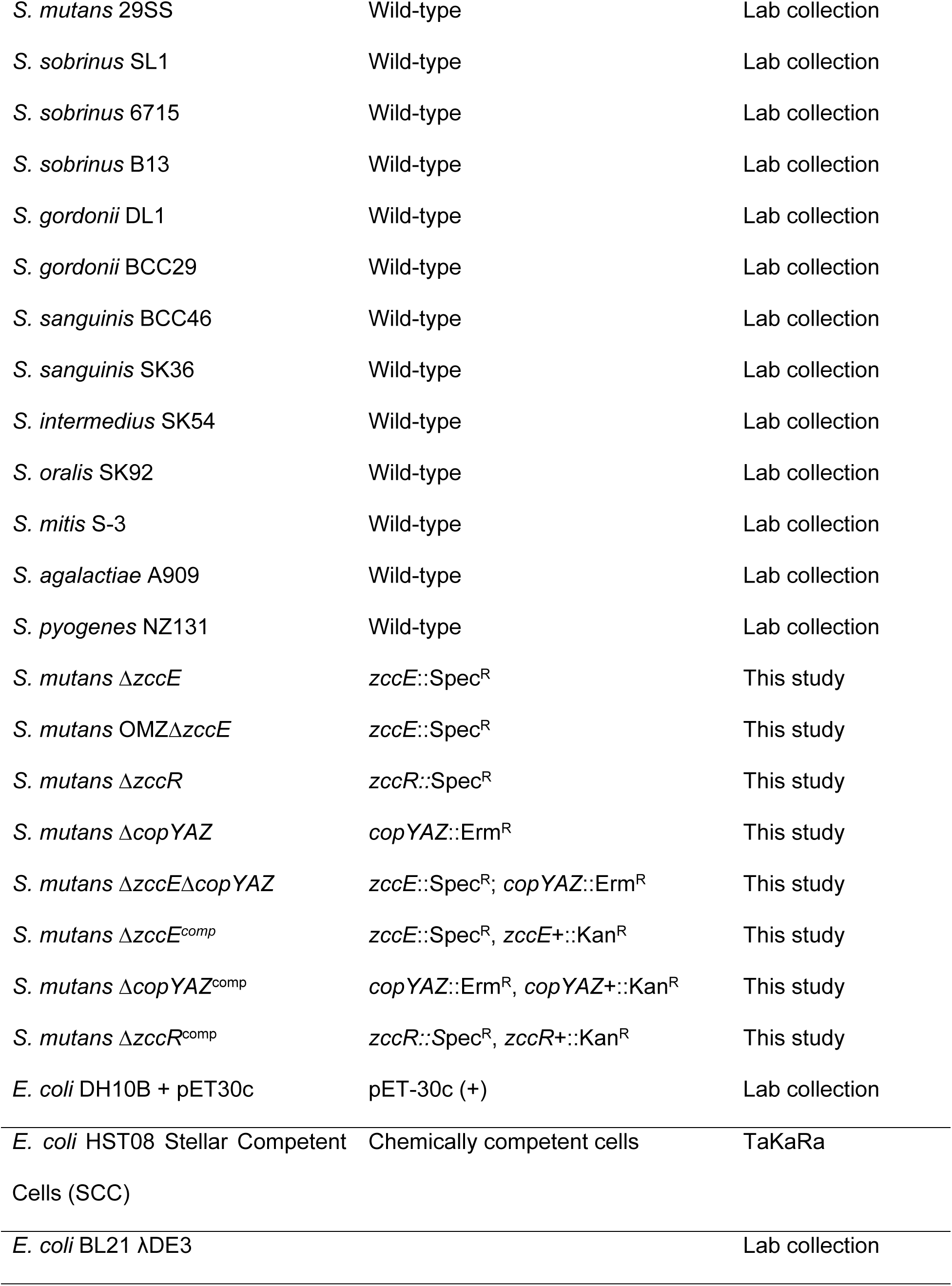
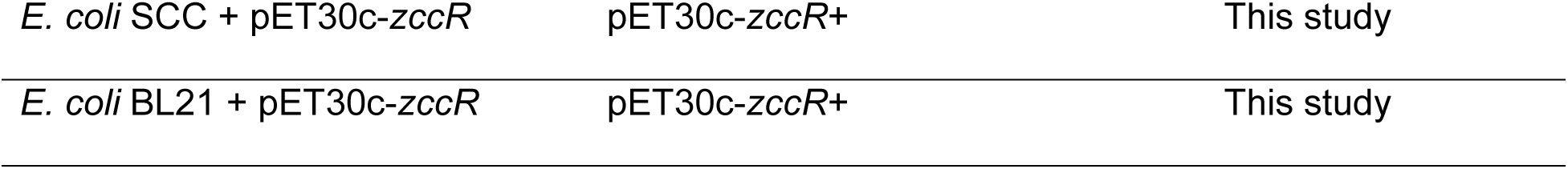
Bacterial strains used in the study.

### Construction of mutant and complemented strains

Strains lacking *zccE*, *zccR*, or the *copYAZ* operon were constructed using a standard PCR ligation mutagenesis approach (65). Briefly, DNA fragments (∼ 1 kb) flanking the target gene were amplified by PCR, digested with restriction enzymes, and ligated to a similarly-digested nonpolar spectinomycin resistance cassette for constructing Δ*zccE* and Δ*zccR* strains and nonpolar erythromycin resistance cassette to obtain the Δ*copYAZ* strain. The ligation mixtures were used to transform *S. mutans* UA159 in the presence of XIP (*co<u>m</u>X*-inducing <u>p</u>eptide) following an established protocol (66). Mutant strains were selected on BHI agar supplemented with the appropriate antibiotic and gene deletions confirmed by PCR amplification and Sanger sequencing. The double Δ*zccE*Δ*copYAZ* mutant was obtained by transforming the Δ*copYAZ* strain with the *zccE*::Spec^R^ region. To generate complemented strains, the full length *zccE, zccR*, or *copYAZ* coding sequences with their native promoters were amplified and cloned into the integration vector pMC340B (67) to yield plasmid pMC340B-*zccE*, pMC340B-*zccR,* and pMC340B-*copYAZ*, respectively. Plasmids were propagated in *E. coli* DH10B and used to transform the *S. mutans* strains Δ*zccE*, Δ*zccR*, or Δ*copYAZ* for integration at the mannitol utilization (*mtl*) locus. All primers used in this study are listed in **Table S3**.

### RNA analysis

Total RNA was isolated from homogenized *S. mutans* cell lysates by acid-phenol chloroform extractions as previously described (50). Total RNA isolated was treated with Ambion DNase I (ThermoFisher) for 30 min at 37°C and further purified using an RNeasy kit (Qiagen), which included a second on-column DNase digestion. Sample quality and quantity were assessed on an Agilent 2100 Bioanalyzer at the University of Florida Interdisciplinary Center for Biotechnology Research (UF-ICBR). For RNA-Seq analysis, samples were prepared as previously described (50). Briefly, 5 µg of the total RNA isolated was subjected to two rounds of mRNA enrichment using a MICROBExpress bacterial mRNA purification kit (Thermo Fisher), and 100 ng of the enriched mRNA used to generate cDNA libraries with unique barcodes (Next UltraII Directional RNA Library Prep kit for Illumina, New England Biolabs). The cDNA libraries were pooled and subjected to RNA-Seq analysis at the UF-ICBR using the Illumina NextSeq 500 platform. Read mapping was performed on a Galaxy server using Map with Bowtie for Illumina and the *S. mutans* UA159 genome (GenBank accession no. NC_004350.2) as a reference. The reads per open reading frame were tabulated with htseq-count. Final comparisons between control and Zn-treated conditions were performed with Degust (http://degust.erc.monash.edu/), with a false-discovery rate (FDR) of 0.05 and 2-fold change cutoff. Quantifications of *zccE* and *gyrA* mRNA were determined by qRT-PCR following an established protocol (68) using the primers listed in **Table S3**. The fold change of *zccE* expression was performed using ΔΔ*C_T_* (where *C_T_* is the threshold cycle) method, with *gyrA* being used as standardization control. One-way ANOVA was performed to verify significance of the qRT-PCR results.

### Metal sensitivity assays

To test sensitivity to Zn in disc diffusion assays, 25 µl of exponentially grown cultures (OD_600_ ∼ 0.5) were spread onto agar plates using a sterile swab and topped with sterile Whatman filter paper discs (6-mm diameter) saturated with 20 μl of a 1 mM ZnSO_4_ solution. The diameter of the zone of growth inhibition was measured after 24 h of incubation at 37°C in 5% CO_2_. The minimum inhibitory concentration (MIC) of ZnSO_4_ was determined by a broth microdilution method using two-fold serial dilutions of ZnSO_4_. Plates were incubated at 37°C in 5% CO_2_ for 16 h and the concentration of ZnSO_4_ at which the absorbance values were 10% of the control condition was determined to be the MIC. For plate titration assays, exponentially grown BHI cultures (OD_600_ ∼ 0.5) were serially diluted and 8 μl of each 10-fold dilution spotted on BHI plates supplemented with selected trace metals. Plates were photographed after 24 h incubation at 37°C in 5% CO_2_ or under anaerobiosis (GasPak jar, BD Biosciences).

### Antagonism assay

The ability of *S. gordonii* to inhibit the growth of *S. mutans* via H_2_O_2_ production was assessed as described previously (50). Briefly, 8 μl of an overnight culture of *S. gordonii* DL1 was spotted on BHI plates with or without Zn (50 μM ZnSO_4_) supplementation and incubated at 37°C in 5% CO_2_. After 24 h incubation, 8 μl of an overnight culture of *S. mutans* UA159 or Δ*zccE* was spotted proximal to the *S. gordonii* spot. To confirm that growth inhibition was due to H_2_O_2_ production, a control condition was included in which 8 μl of 1 mg ml^−1^ catalase solution was spotted on top of the *S. gordonii* spot prior to spotting *S. mutans*.

### ICP-MS analysis

The intracellular Zn and Mn content in parent and mutant strains was determined via inductively <u>c</u>oupled <u>p</u>lasma <u>m</u>ass <u>s</u>pectrometry (ICP-MS) performed at the University of Florida Institute of Food and Agricultural Sciences (UF-IFAS) Analytical Services Laboratories. Briefly, cultures (250 ml) were grown in BHI to mid-exponential phase (OD_600_ 0.4), harvested by centrifugation at 4°C for 15 min at 4,000 rpm, washed in PBS supplemented with 0.2 mM EDTA (to chelate extracellular divalent cations) followed by a second wash in PBS only. The cell pellets were resuspended in 35% (v/v) HNO_3_, heated at 95°C for 1 h before diluted to 3.5% (v/v) HNO_3_ using metal free water and the intracellular Zn and Mn content determined using a 7900 ICP Mass Spectrometer (Agilent). Metal concentrations were normalized to total protein content determined by the bicinchoninic acid (BCA) assay (Pierce).

### ZccR purification

To overexpress and purify a recombinant His-tagged ZccR, the 795-bp *zccR* coding region was amplified by PCR using the primers listed in **Table S3**, and the amplicon cloned onto the expression vector pET30c (Novagen) using the In-fusion HD Cloning Plus kit (TaKaRa) to generate plasmid pET30c-*zccR*. The resultant plasmid was confirmed by Sanger sequencing and transformed into *E. coli* BL21 λDE3. The resulting *E. coli* strain harboring pET30c-*zccR* was grown in a shaking 37°C incubator in Luria-Bertani broth to mid-exponential phase (OD_600_ 0.4). Recombinant His-tagged was ZccR overexpressed by addition of 0.2 mM isopropyl-β-D-1- thiogalactopyranoside (IPTG) (Teknova) and the culture was incubated for additional 3 h. Cell pellets were collected by centrifugation, resuspended in lysis buffer (50 mM NaH_2_PO_4_, 300 mM NaCl, 10 mM imidazole, pH 8) containing lysozyme (1 mg ml^-1^) and sonicated to clarity before centrifugation of cellular debris. The cleared lysate supernatant was mixed with pre-cleared Ni-NTA beads at 4°C for 60 minutes before loaded onto a chromatography column. The packed column was washed with wash buffer (50 mM NaH_2_PO_4_, 300 mM NaCl, 20 mM imidazole), and His-ZccR eluted with elution buffer (50 mM NaH_2_PO_4_, 300 mM NaCl, 250 mM imidazole) according to the QIAexpressionist protocol (Qiagen) for purification of native proteins with histidine tags. The elution fractions were analyzed by 12% SDS-PAGE, and the fractions containing highly pure ZccR pooled and dialyzed against PBS. Aliquots of purified protein were stored at -20°C in 15% glycerol until further use.

### Electrophoretic mobility shift assays

EMSAs were performed with slight modifications from established protocols (50, 69). Briefly, the intergenic region (IGR) between *zccE* and *zccR* was amplified using the primers listed in **Table S3**, and the resulting 160-bp amplicon biotin-labeled using the Biotin 3’ End DNA Labeling kit (ThermoFisher). EMSA reactions were prepared in 20 µl reaction mixtures in binding buffer (10 mM Tris, 50 mM KCl, 1 µg poly dI-dC, 1 mM DTT; 5% glycerol, pH 7.5) containing 20 fmol of labeled probe and 0 to 1.4 µM of purified ZccR. Samples were loaded onto 6% non-denaturing polyacrylamide gels and resolved at 100V for 1h in cold buffer (4°C). Gels were transferred to BrightStar^TM^-Plus positively charged nylon membranes (Thermo-Fisher) and bands visualized using the Chemiluminescent Nucleic Acid Detection Module (Thermo-Fisher) following manufacturer’s protocol. To determine binding specificity, 100X molar excess of either non-labeled specific competitor DNA (*zccE*-*zccR* IGR) or non-specific competitor DNA (206-bp *S. mutans mntH* promoter region (50)) was added to the reaction.

### Oral colonization rat model

A modification of a rat caries model that we have previously followed (38) was used to determine the ability of the Δ*zccE* mutant strain to colonize rats fed a cariogenic diet while also testing the effects of topical Zn treatment on *S. mutans* colonization efficiency. Briefly, specific pathogen-free Sprague-Dawley rat pups were purchased with their dams from Envigo Laboratories and screened upon arrival to ensure an absence of mutans streptococci by plating oral swabbings on mitis salivarius (MS) agar. Prior to infection, pups and dams received 0.8 mg ml^-1^ sulfamethoxazole and 0.16 mg ml^-1^ trimethoprim in the drinking water for 3 days to suppress endogenous flora and facilitate colonization by *S. mutans*. After 4 days of washout during which antibiotic-free water was provided, dams and pups aged 18 days were orally infected for four consecutive days with actively growing *S. mutans* UA159 or Δ*zccE* cultures by means of cotton swab. At the time of infection, the regular chow diet was replaced by a 12% sucrose cariogenic powdered diet (ENVIGO diet TD.190707). During the infection period, the animals were provided with 5% (wt/vol) sterile sucrose-water *ad libitum*, then fresh water for the remainder of the study. On the final day of infection, pups were weaned and randomly placed into experimental groups. Topical treatment with ZnSO_4_ or saline solutions commenced after the fourth and final day of bacterial infection and lasted 10 consecutive days. Test (60 mM or 150 mM ZnSO_4_) or control (saline) treatments were topically administered to the rat teeth with a camel’s hairbrush twice a day with a 6 h interval between treatments. At the end of the treatment period, animals were euthanized by CO_2_ asphyxiation, and the lower jaws removed for bacterial burden determination. Jaw sonicates were subjected to 10-fold serial dilutions and plated on MS agar (to count *S. mutans*) and 5% sheep blood agar (to count total flora). The number of *S. mutans* recovered from the animals was expressed as CFU ml-1 of jaw sonicate, and *S. mutans* colonies counted on MS agar divided by the total CFU on blood agar to determine the percentage of *S. mutans* colonies recovered over the total flora. This study was reviewed and approved by the University of Florida Institutional Animal Care and Use Committee (protocol # 201810421).

## Ethics statement

Animal procedure for rat colorizations was approved by the University of Florida Institutional Animal Care and Use Committee (protocol # 201810421). All animal care was consistent with the Guide for the Care and Use of Laboratory Animals from the National Research Council and the USDA Animal Care Resource Guide.

## Data availability

Gene expression data have been deposited in the NCBI Gene Expression Omnibus (GEO) database (https://www.ncbi.nlm.nih.gov/geo) under GEO Series accession number GSE166993.

## Acknowledgements

This study was supported by NIH-NIDCR award R01 DE019783 to J.A.L. A.M.P was also supported by T90 DE021990.

## Supporting information

Fig S1. Growth curves of *S. mutans* UA159 in the chemically-defined FMC medium spiked with 0, 2, 4 or 6 mM ZnSO_4_ upon reaching mid-logarithmic phase (OD_600_ ∼ 0.3).

Fig S2. Growth of *S. mutans* OMZ175 and Δ*zccE* derivative in BHI with or without 1 mM ZnSO_4_ supplementation.

Fig S3. Plate titration (spot test) of *S. mutans* UA159, Δ*zccE*, Δ*copYAZ* and Δcop*YAZ*Δ*zccE* strains on BHI plates supplemented with increasing concentration of CuSO_4_ and incubated for 24 h at 37°C under anaerobic conditions (Gaspack). Images are representative of at least 3 independent experiments.

Fig S4. Growth curves of UA159, Δ*zccE*, Δ*zccR* and respective complemented strains in BHI with or without Zn supplementation (150 µM ZnSO_4_).

Fig S5. Growth of *S. mutans* UA159 and *ΔzccE* strains with Zn and Mn. (A) Growth in BHI medium with supplementation of Zn and Mn in different ratios. (B) Growth inhibition zones for *S. mutans* UA159 and Δ*zccE* strains grown on BHI agar containing 250 µM Mn with or without Zn supplementation and exposed to filter paper discs saturated with 0.25% H_2_O_2_.

Fig S6. Determination of Zn MIC for UA159 in BHI media in presence of Na-orthovanadate. *S. mutans* UA159 was grown in BHI with increasing amount of Na-orthovanadate and Zn simultaneously to determine Zn MIC.

## References

1. Chandrangsu P, Rensing C, Helmann JD. Metal homeostasis and resistance in bacteria. Nat Rev Microbiol. 2017;15(6):338–50.

2. Palmer LD, Skaar EP. Transition Metals and Virulence in Bacteria. Annu Rev Genet. 2016;50:67–91.

3. Sheldon JR, Skaar EP. Metals as phagocyte antimicrobial effectors. Curr Opin Immunol. 2019;60:1–9.

4. Monteith AJ, Skaar EP. The impact of metal availability on immune function during infection. Trends Endocrinol Metab. 2021;32(11):916–28.

5. Cassat JE, Skaar EP. Iron in infection and immunity. Cell Host Microbe. 2013;13(5):509–19.

6. Zygiel EM, Nolan EM. Transition Metal Sequestration by the Host-Defense Protein Calprotectin. Annu Rev Biochem. 2018;87:621–43.

7. Djoko KY, Ong CL, Walker MJ, McEwan AG. The Role of Copper and Zinc Toxicity in Innate Immune Defense against Bacterial Pathogens. J Biol Chem. 2015;290(31):18954–61.

8. Lonergan ZR, Skaar EP. Nutrient Zinc at the Host-Pathogen Interface. Trends Biochem Sci. 2019;44(12):1041–56.

9. Ong CL, Gillen CM, Barnett TC, Walker MJ, McEwan AG. An antimicrobial role for zinc in innate immune defense against group A streptococcus. J Infect Dis. 2014;209(10):1500–8.

10. Makthal N, Kumaraswami M. Zinc’ing it out: zinc homeostasis mechanisms and their impact on the pathogenesis of human pathogen group A streptococcus. Metallomics. 2017;9(12):1693–702.

11. Hantke K. Bacterial zinc transporters and regulators. Biometals. 2001;14(3-4):239–49.

12. Moore CM, Helmann JD. Metal ion homeostasis in Bacillus subtilis. Curr Opin Microbiol. 2005;8(2):188–95.

13. Beard SJ, Hashim R, Membrillo-Hernandez J, Hughes MN, Poole RK. Zinc(II) tolerance in Escherichia coli K-12: evidence that the zntA gene (o732) encodes a cation transport ATPase. Mol Microbiol. 1997;25(5):883–91.

14. Gaballa A, Helmann JD. Bacillus subtilis CPx-type ATPases: characterization of Cd, Zn, Co and Cu efflux systems. Biometals. 2003;16(4):497–505.

15. Rensing C, Mitra B, Rosen BP. The zntA gene of Escherichia coli encodes a Zn(II)- translocating P-type ATPase. Proc Natl Acad Sci U S A. 1997;94(26):14326–31.

16. Martin JE, Edmonds KA, Bruce KE, Campanello GC, Eijkelkamp BA, Brazel EB, et al. The zinc efflux activator SczA protects Streptococcus pneumoniae serotype 2 D39 from intracellular zinc toxicity. Mol Microbiol. 2017;104(4):636–51.

17. Martin JE, Giedroc DP. Functional Determinants of Metal Ion Transport and Selectivity in Paralogous Cation Diffusion Facilitator Transporters CzcD and MntE in Streptococcus pneumoniae. J Bacteriol. 2016;198(7):1066–76.

18. Jacobsen FE, Kazmierczak KM, Lisher JP, Winkler ME, Giedroc DP. Interplay between manganese and zinc homeostasis in the human pathogen Streptococcus pneumoniae. Metallomics. 2011;3(1):38–41.

19. Kloosterman TG, van der Kooi-Pol MM, Bijlsma JJ, Kuipers OP. The novel transcriptional regulator SczA mediates protection against Zn2+ stress by activation of the Zn2+-resistance gene czcD in Streptococcus pneumoniae. Mol Microbiol. 2007;65(4):1049–63.

20. Anton A, Grosse C, Reissmann J, Pribyl T, Nies DH. CzcD is a heavy metal ion transporter involved in regulation of heavy metal resistance in Ralstonia sp. strain CH34. J Bacteriol. 1999;181(22):6876–81.

21. Ong CL, Walker MJ, McEwan AG. Zinc disrupts central carbon metabolism and capsule biosynthesis in Streptococcus pyogenes. Sci Rep. 2015;5:10799.

22. Sullivan MJ, Goh KGK, Ulett GC. Cellular Management of Zinc in Group B Streptococcus Supports Bacterial Resistance against Metal Intoxication and Promotes Disseminated Infection. mSphere. 2021;6(3).

23. Selwitz RH, Ismail AI, Pitts NB. Dental caries. Lancet. 2007;369(9555):51-9.

24. The Lancet Child Adolescent H. Oral health: oft overlooked. Lancet Child Adolesc Health. 2019;3(10):663.

25. Pant S, Patel NJ, Deshmukh A, Golwala H, Patel N, Badheka A, et al. Trends in infective endocarditis incidence, microbiology, and valve replacement in the United States from 2000 to 2011. J Am Coll Cardiol. 2015;65(19):2070–6.

26. Chamat-Hedemand S, Dahl A, Ostergaard L, Arpi M, Fosbol E, Boel J, et al. Prevalence of Infective Endocarditis in Streptococcal Bloodstream Infections Is Dependent on Streptococcal Species. Circulation. 2020;142(8):720–30.

27. Lynch RJ. Zinc in the mouth, its interactions with dental enamel and possible effects on caries; a review of the literature. Int Dent J. 2011;61 Suppl 3:46–54.

28. Uwitonze AM, Ojeh N, Murererehe J, Atfi A, Razzaque MS. Zinc Adequacy Is Essential for the Maintenance of Optimal Oral Health. Nutrients. 2020;12(4).

29. Giertsen E. Effects of mouthrinses with triclosan, zinc ions, copolymer, and sodium lauryl sulphate combined with fluoride on acid formation by dental plaque in vivo. Caries Res. 2004;38(5):430–5.

30. Nossek H, Dobl P. [The effect of zinc chloride mouthwashes on caries-inducing plaque streptococci. 3. The antibacterial effect of zinc chloride on the species Str. mutans, Str. sanguis and Str. salivarius in dental plaque]. Zahn Mund Kieferheilkd Zentralbl. 1990;78(6):501–5.

31. Compton FH, Beagrie GS. Inhibitory effect of benzethonium and zinc chloride mouthrinses on human dental plaque and gingivitis. J Clin Periodontol. 1975;2(1):33–43.

32. Fischman SL, Picozzi A, Cancro LP, Pader M. The inhibition of plaque in humans by two experimental oral rinses. J Periodontol. 1973;44(2):100–2.

33. Bates DG, Navia JM. Chemotherapeutic effect of zinc on streptococcus mutans and rat dental caries. Arch Oral Biol. 1979;24(10-11):799–805.

34. Phan TN, Buckner T, Sheng J, Baldeck JD, Marquis RE. Physiologic actions of zinc related to inhibition of acid and alkali production by oral streptococci in suspensions and biofilms. Oral Microbiol Immunol. 2004;19(1):31–8.

35. He G, Pearce EI, Sissons CH. Inhibitory effect of ZnCl(2) on glycolysis in human oral microbes. Arch Oral Biol. 2002;47(2):117–29.

36. Izaguirre-Fernandez EJ, Eisenberg AD, Curzon ME. Interactions of zinc with fluoride on growth, glycolysis and survival of Streptococcus mutans GS-5. Caries Res. 1989;23(1):18–25.

37. Koo H, Sheng J, Nguyen PT, Marquis RE. Co-operative inhibition by fluoride and zinc of glucosyl transferase production and polysaccharide synthesis by mutans streptococci in suspension cultures and biofilms. FEMS Microbiol Lett. 2006;254(1):134–40.

38. Ganguly T, Peterson AM, Kajfasz JK, Abranches J, Lemos JA. Zinc import mediated by AdcABC is critical for colonization of the dental biofilm by Streptococcus mutans in an animal model. Mol Oral Microbiol. 2021;36(3):214–24.

39. Tatusov RL, Galperin MY, Natale DA, Koonin EV. The COG database: a tool for genome-scale analysis of protein functions and evolution. Nucleic Acids Res. 2000;28(1):33–6.

40. Singh K, Senadheera DB, Levesque CM, Cvitkovitch DG. The copYAZ Operon Functions in Copper Efflux, Biofilm Formation, Genetic Transformation, and Stress Tolerance in Streptococcus mutans. J Bacteriol. 2015;197(15):2545–57.

41. Young CA, Gordon LD, Fang Z, Holder RC, Reid SD. Copper Tolerance and Characterization of a Copper-Responsive Operon, copYAZ, in an M1T1 Clinical Strain of Streptococcus pyogenes. J Bacteriol. 2015;197(15):2580–92.

42. Fu Y, Tsui HC, Bruce KE, Sham LT, Higgins KA, Lisher JP, et al. A new structural paradigm in copper resistance in Streptococcus pneumoniae. Nat Chem Biol. 2013;9(3):177–83.

43. Fan B, Rosen BP. Biochemical characterization of CopA, the Escherichia coli Cu(I)- translocating P-type ATPase. J Biol Chem. 2002;277(49):46987–92.

44. Silver S. Bacterial resistances to toxic metal ions--a review. Gene. 1996;179(1):9–19.

45. Vats N, Lee SF. Characterization of a copper-transport operon, copYAZ, from Streptococcus mutans. Microbiology. 2001;147(Pt 3):653–62.

46. Shafeeq S, Yesilkaya H, Kloosterman TG, Narayanan G, Wandel M, Andrew PW, et al. The cop operon is required for copper homeostasis and contributes to virulence in Streptococcus pneumoniae. Mol Microbiol. 2011;81(5):1255–70.

47. Cooksey DA. Molecular mechanisms of copper resistance and accumulation in bacteria. FEMS Microbiol Rev. 1994;14(4):381–6.

48. Rae TD, Schmidt PJ, Pufahl RA, Culotta VC, O’Halloran TV. Undetectable intracellular free copper: the requirement of a copper chaperone for superoxide dismutase. Science. 1999;284(5415):805–8.

49. Neubert MJ, Dahlmann EA, Ambrose A, Johnson MDL. Copper Chaperone CupA and Zinc Control CopY Regulation of the Pneumococcal cop Operon. mSphere. 2017;2(5).

50. Kajfasz JK, Katrak C, Ganguly T, Vargas J, Wright L, Peters ZT, et al. Manganese Uptake, Mediated by SloABC and MntH, Is Essential for the Fitness of Streptococcus mutans. mSphere. 2020;5(1).

51. Bosma EF, Rau MH, van Gijtenbeek LA, Siedler S. Regulation and distinct physiological roles of manganese in bacteria. FEMS Microbiol Rev. 2021;45(6).

52. Ortiz de Orue Lucana D, Wedderhoff I, Groves MR. ROS-Mediated Signalling in Bacteria: Zinc-Containing Cys-X-X-Cys Redox Centres and Iron-Based Oxidative Stress. J Signal Transduct. 2012;2012:605905.

53. Stephen KW, Creanor SL, Russell JI, Burchell CK, Huntington E, Downie CF. A 3-year oral health dose-response study of sodium monofluorophosphate dentifrices with and without zinc citrate: anti-caries results. Community Dent Oral Epidemiol. 1988;16(6):321–5.

54. Parkinson CR, Burnett GR, Creeth JE, Lynch RJM, Budhawant C, Lippert F, et al. Effect of phytate and zinc ions on fluoride toothpaste efficacy using an in situ caries model. J Dent. 2018;73:24–31.

55. Pace NJ, Weerapana E. Zinc-binding cysteines: diverse functions and structural motifs. Biomolecules. 2014;4(2):419–34.

56. O’Brien J, Pastora A, Stoner A, Spatafora G. The S. mutans mntE gene encodes a manganese efflux transporter. Mol Oral Microbiol. 2020;35(3):129–40.

57. Lutsenko S, Kaplan JH. Organization of P-type ATPases: significance of structural diversity. Biochemistry. 1995;34(48):15607–13.

58. Solioz M, Vulpe C. CPx-type ATPases: a class of P-type ATPases that pump heavy metals. Trends Biochem Sci. 1996;21(7):237–41.

59. Dutta SJ, Liu J, Stemmler AJ, Mitra B. Conservative and nonconservative mutations of the transmembrane CPC motif in ZntA: effect on metal selectivity and activity. Biochemistry. 2007;46(12):3692–703.

60. Eijkelkamp BA, Morey JR, Ween MP, Ong CL, McEwan AG, Paton JC, et al. Extracellular zinc competitively inhibits manganese uptake and compromises oxidative stress management in Streptococcus pneumoniae. PLoS One. 2014;9(2):e89427.

61. McDevitt CA, Ogunniyi AD, Valkov E, Lawrence MC, Kobe B, McEwan AG, et al. A molecular mechanism for bacterial susceptibility to zinc. PLoS Pathog. 2011;7(11):e1002357.

62. Spatafora G, Corbett J, Cornacchione L, Daly W, Galan D, Wysota M, et al. Interactions of the Metalloregulatory Protein SloR from Streptococcus mutans with Its Metal Ion Effectors and DNA Binding Site. J Bacteriol. 2015;197(22):3601–15.

63. Magalhaes PP, Paulino TP, Thedei G, Jr., Larson RE, Ciancaglini P. A 100 kDa vanadate and lanzoprazole-sensitive ATPase from Streptococcus mutans membrane. Arch Oral Biol. 2003;48(12):815–24.

64. Terleckyj B, Willett NP, Shockman GD. Growth of several cariogenic strains of oral streptococci in a chemically defined medium. Infect Immun. 1975;11(4):649–55.

65. Lau PC, Sung CK, Lee JH, Morrison DA, Cvitkovitch DG. PCR ligation mutagenesis in transformable streptococci: application and efficiency. J Microbiol Methods. 2002;49(2):193–205.

66. Morrison DA, Khan R, Junges R, Amdal HA, Petersen FC. Genome editing by natural genetic transformation in Streptococcus mutans. J Microbiol Methods. 2015;119:134–41.

67. Chen PM, Chen YY, Yu SL, Sher S, Lai CH, Chia JS. Role of GlnR in acid-mediated repression of genes encoding proteins involved in glutamine and glutamate metabolism in Streptococcus mutans. Appl Environ Microbiol. 2010;76(8):2478–86.

68. Kajfasz JK, Ganguly T, Hardin EL, Abranches J, Lemos JA. Transcriptome responses of Streptococcus mutans to peroxide stress: identification of novel antioxidant pathways regulated by Spx. Sci Rep. 2017;7(1):16018.

69. Rolerson E, Swick A, Newlon L, Palmer C, Pan Y, Keeshan B, et al. The SloR/Dlg metalloregulator modulates Streptococcus mutans virulence gene expression. J Bacteriol. 2006;188(14):5033–44.

